# PCID2 dysregulates transcription and viral RNA processing to promote HIV-1 latency

**DOI:** 10.1101/2023.09.21.558802

**Authors:** Raquel Crespo, Enrico Ne, Julian Reinders, Jenny I. J. Meier, Chengcheng Li, Sanne Jansen, Alicja Górska, Selin Koçer, Tsung Wai Kan, Wouter Doff, Dick Dekkers, Jeroen Demmers, Robert-Jan Palstra, Shringar Rao, Tokameh Mahmoudi

## Abstract

HIV-1 latency results from tightly regulated molecular processes that act at distinct steps of HIV-1 gene expression. To elucidate the molecular players that govern latency, we previously performed a dCas9-chromatin immunoprecipitation coupled with mass spectrometry (Catchet-MS) and identified the interactome of the latent HIV-1 LTR. Here we characterize the Catchet-MS-identified PCI domain-containing 2 (PCID2) protein, a component of the TREX2 complex, to play a dual role in promoting HIV-1 latency by enforcing both transcriptional repression and post-transcriptional blocks to HIV-1 gene expression. PCID2 bound the latent HIV-1 LTR and repressed transcription initiation during latency. Depletion of PCID2 remodelled the chromatin landscape at the HIV-1 promoter and resulted in transcriptional activation and reversal of latency. Immunoprecipitation coupled to Mass Spectrometry identified PCID2-interacting proteins to include members of the spliceosome, including negative viral RNA (vRNA) alternative splicing regulators, and PCID2 depletion resulted in over-splicing of intron-containing vRNA and misregulated expression of vRNA splice variants. We demonstrate that MCM3AP and DSS1, two other RNA-binding TREX2 complex subunits that comprise the dock of the complex also inhibit transcription initiation and viral RNA alternative splicing during latency and similarly to PCID2 function as prominent latency associated repressors of HIV-1 gene expression. Thus, PCID2 is a novel HIV-1 latency-promoting factor, which in context of the TREX2 sub-complex PCID2-DSS1-MCM3AP blocks transcription and dysregulates vRNA processing.

## Introduction

The persistence of latent HIV-1 viral reservoirs in memory CD4+ T cells that elude elimination by the immune system and cannot be targeted by antiretroviral therapy is considered the biggest obstacle toward an HIV-1 cure ^1–3^. Latency is established upon integration of the viral genome into the host cell genome and is defined by the absence of viral production ^3^. The establishment and maintenance of viral latency is a layered multifactorial process that is enforced by distinct host cell pathways acting on chromatin regulation, transcription initiation and elongation, and post-transcriptional mechanisms regulating RNA metabolism and viral protein production ^4,5^.

Transcription of the HIV-1 provirus is directed by its promoter in the 5’ long terminal repeat (LTR) whose activity is controlled by the chromatin landscape and relies on the availability of viral protein Tat and host cell transcription (co-)factors ^5–8^. During latency, the viral protein Tat is absent, which causes RNA Pol II to pause at the TAR RNA element (+70) prohibiting transcription elongation ^9^. Further blocks to HIV-1 gene expression occur at a post-transcriptional level, including inhibition of viral RNA (vRNA) splicing, nucleocytoplasmic export, translation and trafficking (reviewed in ^4^). The contribution of these RNA processing mechanisms to the overall repression of HIV-1 gene expression has recently been characterized in both in vitro models of HIV-1 latency and CD4+ T cells obtained from people living with HIV-1 (PLWH) ^10–14^, where it has been shown that blocks to HIV-1 gene expression occur largely at post-transcriptional levels. More recently, several unbiased and candidate studies, including our own, have successfully identified co– and post-transcriptional regulators of HIV-1 latency that control vRNA metabolism and either promote or inhibit viral gene expression ^12,15,16^.

In a previous study, we performed an HIV-1 LTR locus-specific dCas9 chromatin immunoprecipitation coupled to mass spectrometry (Catchet-MS) in latent and active cells and identified putative LTR-bound transcriptional regulators of HIV-1 latency ^16^. Interestingly, while a substantial number of latency-associated factors identified were, as expected, DNA associated, we also identified many putative RNA-binding latent HIV-1 promoter enriched regulators involved in RNA metabolism, including CRNKL1, recently described to post-transcriptionally regulate HIV-1 gene expression ^17^, and other known regulators of HIV-1 gene expression such as DDX24 ^18^.

Catchet-MS also identified the host protein PCI domain containing 2 (PCID2) to be enriched at the HIV-1 promoter in the latent state. PCID2 (human homologue of yeast protein Thp1) is mainly described in literature as part of the multi-subunit Transcription and Export Complex 2 (TREX2), composed of PCID2, GANP, DSS1, ENY2 and CETN2/3 subunits, which links transcription with mRNA export by binding to nascent RNA containing messenger ribonucleoprotein (mRNP) complexes and mediating their transport from transcription sites to the nuclear pore for subsequent export ^19–23^. Outside of the TREX2 complex, other functions have been ascribed to PCID2, including roles in genome instability and preventing R-Loop formation ^24,25^, in protein stability ^26,27^ and transcription regulation of lymphoid commitment genes ^28^.

Here, we demonstrate that PCID2 plays a dual role during latency, acting both at the level of HIV-1 transcription initiation and viral RNA processing steps, resulting in repression of HIV-1 gene expression. Chromatin immunoprecipitation experiments indicate that, while enriched at the latent HIV-1 LTR, PCID2 is removed upon transcriptional activation, and acts as a repressor of HIV-1 gene expression by inhibiting LTR-driven transcription initiation. Downregulation of PCID2 leads to overall de-repression of the HIV-1 LTR locus as indicated by removal or deposition of distinct chromatin marks concomitant with recruitment of activating transcription factors to the promoter. In a transcription-independent mechanism, PCID2 also promotes latency by blocking post-transcriptional steps of HIV-1 gene expression. Immunoprecipitation of PCID2 coupled to mass spectrometry identified PCID2 interaction partners involved in RNA splicing, export, stability and translation. Interestingly, downregulation of PCID2 led to the over-splicing of HIV-1 vRNAs and mis-regulated expression of HIV-1 RNA splice variants, indicating that PCID2 promotes HIV-1 latency by inhibiting alternative HIV-1 RNA splicing. We find that TREX2 subunits MCM3AP and DSS1 also enforce viral latency by inhibiting transcription initiation and viral RNA splicing, indicating that PCID2-mediated repression of HIV-1 gene expression occurs in context of the RNA-binding PCID2-DSS1-MCM3AP sub-complex and providing a putative role for the TREX2 complex. Consistent with its function within the TREX2 complex, PCID2 regulates nucleocytoplasmic export of completely spliced viral RNA species. Thus, PCID2-containing complexes play a dual role to promote HIV-1 latency by blocking both transcription initiation and vRNA splicing, and facilitates export of completely spliced vRNA species.

## Results

### PCID2 is an HIV-1 LTR-bound latency promoting factor

In a previous study ^16^, we identified putative latent promoter-bound regulators of HIV-1 gene expression by dCas9-targeted chromatin immunoprecipitation coupled to mass spectrometry (Catchet-MS) (Figure 1A). Because of differential enrichment at the latent HIV-1 and inactive LTR, we hypothesized that these factors may contribute to the maintenance of HIV-1 latency as putative repressors of HIV-1 gene expression. Interestingly, more than half of the latent HIV-1 LTR-bound proteins identified were categorized to be RNA binding and/or involved in host RNA metabolism (Figure 1A). Amongst these, endogenous PCI-domain containing protein 2 (PCID2) was found by the Catchet-MS pipeline to be preferentially associated with the repressed HIV-1 promoter in latent J-Lat 11.1 cells.

**Figure 1.**
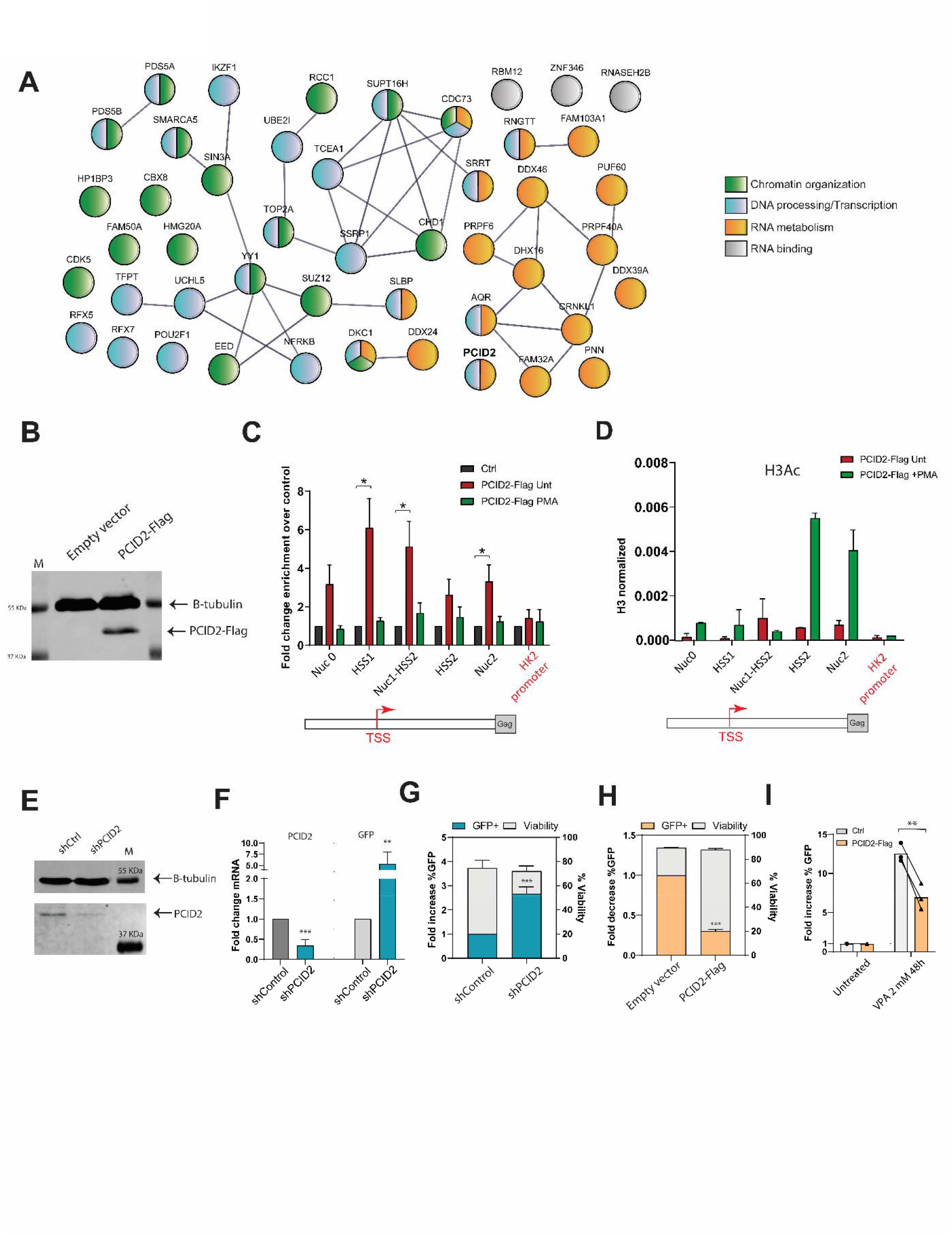
PCID2 is enriched at the latent HIV-1 LTR and is a repressor of HIV-1 gene expression during latency. **A**. STRING network of proteins identified to be enriched in the latent HIV-1 LTR by the Catchet-MS pipeline as published in Ne, Crespo et al. 2022 Nuc Acid Res. Proteins are colored based on GO analysis enrichment as indicated in the figure. Grey lines represent known interaction (experiments) as reported in the STRING database. **B**. Western blot of PCID2-Flag in control or stably PCID2-Flag expressing J-Lat 11.1. Beta tubulin was used as loading control. **C**. Enrichment of PCID2-Flag at the HIV-1 LTR expressed as fold enrichment in control and untreated and PMA treated PCID2-Flag expressing J-Lat 11.1 lines as assessed by chromatin immunoprecipitation (ChIP) coupled with quantitative PCR. Primers spanning across the HIV-1 promoter were used to assess relative enrichment of PCID2-Flag at sequential regions of the LTR. HK2 promoter was used as a control genomic region. Relative enrichment was normalized to Input. Mean and SD correspond to 4 ChIP replicates with independent chromatin preparations for control, untreated, and PMA treated cells. **D**. ChIP-qPCR analysis of acetylated Histone 3 (F) enrichment at the HIV-1 promoter in untreated and PMA treated stably PCID2-Flag expressing J-Lat 11.1 lines. H3Ac enrichment at the HIV-1 LTR is represented as relative enrichment normalized to Total H3 as shown in Supplementary Figure 1 C. ChIP-qPCR corresponds to one chromatin preparation, bars and error lines represent, respectively, mean and SD of technical duplos. **E**. Western blot analysis of PCID2 in control and PCID2 knockdown J-Lat 11.1. Beta-tubulin was used as a loading control. Cells were infected with a VSV-pseudotyped lentivirus containing a scramble shRNA (shControl) or a PCID2 mRNA-targeting shRNA (shPCID2). **F**. Gene expression analysis of shRNA-mediated knockdown of PCID2 and GFP mRNA fold induction in shPCID2 cells relative to shControl and normalized to cyclophilin A. Bars represent mean of 5 independent shRNA-mediated knockdown experiments and error lines represent SEM (n=5). Statistical significance was determined by t-test; **p<0.01, ***p<0.001. **G**. Fold increase in the % of GFP (left y axes) and viability (right y axes) in shControl or shPCID2 J-Lat 11.1 cells as measured by flow cytometry. Bars represent mean of 5 independent shRNA-mediated knockdown experiments and error lines represent SEM. Statistical significance was determined by t-test; ***p<0.001. **H**. Fold decrease in the % of GFP (left y axes) and viability (right y axes) in control or transiently PCID2-Flag overexpressing J-Lat 11.1 cells as measured by flow cytometry. Bars represent mean of 3 collections and error lines represent SEM. Statistical significance was determined by t-test; ***p<0.001. **I**. Fold change in % of GFP in control or PCID2-Flag overexpressing J-Lat 11.1 after a 48 hour treatment of with latency reversing agent valproic acid (VPA; 2 mM) as measured by flow cytometry. Fold increase in % GFP was assessed by normalization to untreated control. Bars represent mean. Statistical significance was determined by t-test; **p<0.01.

We first set out to confirm whether PCID2, is physically bound to the HIV-1 promoter in the latent state. For this, we generated stably transfected J-Lat 11.1 cells which express PCID2-Flag (F-PCID2 J-Lat 11.1), given lack of availability of specific antibodies to purify endogenous PCID2. We validated the exogenous expression of PCID2 by Western blot analysis (Figure 1B), and by RT-PCR confirmed expression levels comparable to endogenous PCID2 (Supplementary Figure 1A). Consistent with Catchet-MS, which identified endogenous PCID2 enriched at the latent HIV-1 LTR, chromatin immunoprecipitation (ChIP) using anti-FLAG M2 beads coupled to qPCR with primers spanning the HIV-1 LTR showed that PCID2 is recruited to the HIV-1 LTR during latency (Figure 1C, Supplementary Figure 1B-C) and enriched at the transcription start site (TSS). We also performed ChIP-qPCR in PCID2-Flag expressing cells treated with the mitogen/PKC agonist phorbol 12-myristate 13-acetate (PMA), that causes strong general transcriptional activation in lymphoid cells (Figure 1C). Upon PMA treatment and transcriptional activation, PCID2 is removed from the HIV-1 promoter (Figure 1C), consistent with its putative role as a repressor and its loss from the LTR in PMA-treated J-Lat 11.1, also observed in Catchet-MS. As expected, PMA treatment of PCID2-Flag expressing cells resulted in an increase in acetylated Histone 3 (Figure 1D, Supplementary Figure 1D-F), indicative of an open chromatin state and active transcription.

To investigate the role of PCID2 during HIV-1 latency, we depleted endogenous PCID2 in J-Lat 11.1 cells with short hairpin RNAs targeting PCID2 mRNA (shPCID2) or scramble control (shCtrl) (Figure 1E-F) and observed a significant increase both in the levels of GFP mRNA and the percentage of cells expressing GFP protein as measured by flow cytometry (Figure 1F-G). To rule out potential contribution of clonal effects, we also depleted PCID2 in J-Lat full length clone 10.6 (Supplementary Figure 2A-B) and LTR-Tat-GFP J-Lat clone A2 (Supplementary Figure 2C-D) and confirmed the presence of significant latency reversal upon PCID2 knockdown. Consistent with these results, transient overexpression of PCID2 decreased the percentage of GFP expressing cells (Figure 1H) and prevented viral reactivation in J-Lat 11.1 cells treated with the latency reversing agent valproic acid (HDAC inhibitor) (Figure 1H). To further validate that PCID2 negatively regulates HIV-1 gene expression, we transiently co-transfected uninfected Jurkat cells with a plasmid containing an env deficient full-length HIV-1 genome and a luciferase reporter replacing the nef gene (pNL4.3.Luc.R-E-) together with either an empty pBud-control or PCID2 expressing pBud-PCID2-Flag plasmid (Supplementary Figure 3A). We observed a significant dose-dependent decrease in luciferase activity in PCID2 transiently overexpressing cells as compared to control cells demonstrating that PCID2 prohibits HIV-1 gene expression in a dose-dependent manner (Supplementary Figure 3B).

Collectively, our results show that PCID2 is present at the HIV-1 LTR during viral latency, is removed upon transcriptional activation, and acts as a repressor of HIV-1 gene expression during HIV-1 latency.

### PCID2 promotes HIV-1 latency by enforcing blocks at transcription initiation

Of the functions currently described for PCID2, two stand out as particularly relevant for its role in HIV latency: first is its function in eukaryotic RNA metabolism in context of the TREX2 complex ^19,20,22^, and second its distinct role in the transcriptional regulation of lymphoid lineage commitment genes ^28^. Because PCID2 differentially binds the latent LTR, we first aimed to determine whether the presence of PCID2 at the HIV-1 promoter mediates its role in repressing HIV-1 gene expression at a transcriptional level. To test this, we performed a modified version of the transcriptional profiling qPCR assay described by Telwatte et al. ^10^ in control and PCID2-knockdown cells using primers specific for amplification of three distinct regions in the LTR: Read through region before the TSS (result of read through transcription from the gene where the proviral genome is integrated), TAR RNA region (region after TSS where RNA Pol II pauses) and U5 UTR region (region indicative of elongated transcripts) (Figure 2A). We calculated the ratio of the relative abundance of different HIV-1 RNA amplicons to assess the relative release of blocks at transcription initiation (TAR/Read Through) and transcription elongation (UTR/TAR). Upon PCID2 knockdown, we observed a significant increase in the TAR/Read Through ratio, but not the UTR/TAR ratio (transcriptional elongation) compared with shControl lines (Figure 2B), indicating a release of a block at transcription initiation, but not transcription elongation, and hence suggesting a role for PCID2 in blocking HIV-1 LTR transcription initiation.

**Figure 2.**
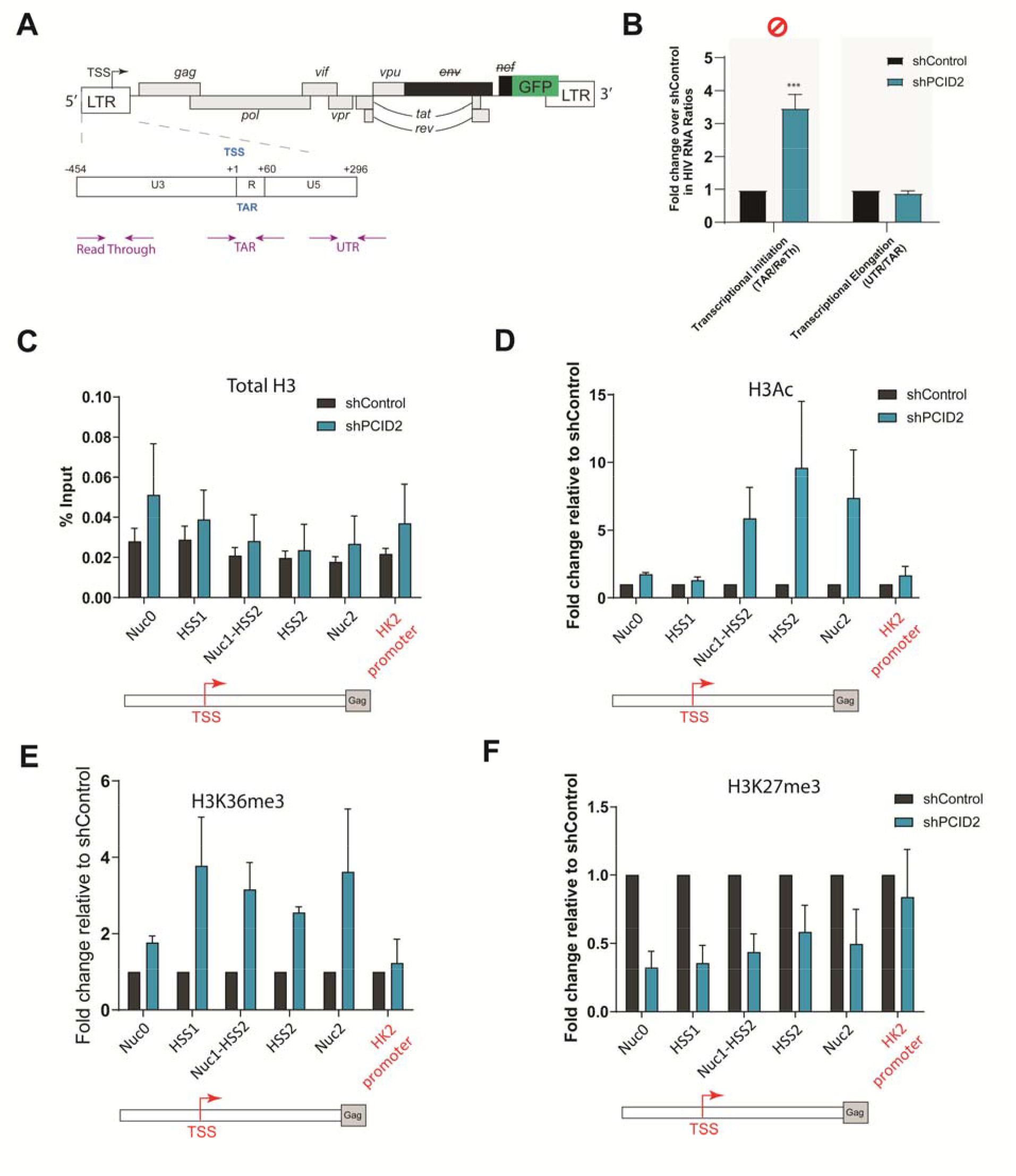
PCID2 is a repressor of HIV-1 transcription initiation during viral latency. **A**. Schematic figure of the HIV-1 genome in J-Lat 11.1. Arrows represent primer location and amplicons across the HIV-1 genome used in the transcriptional profiling assay. **B**. Transcriptional profiling assay of shControl and shPCID2 J-Lat 11.1. Gene expression blocks at transcriptional initiation, elongation, and post-transcriptional steps are assessed by calculating the ratio of the relative abundance of HIV-1 RNA species as shown in the figure. Data is presented as fold change in HIV-1 RNA ratios in shPCID2 relative to shControl. Bars represent mean of 5 independent shRNA-mediated knockdown experiments and error lines represent SEM. Statistical significance was determined by t-test; *p<0.05, ***p<0.001. **C-F**. ChIP-qPCR analysis of total Histone 3 (**C**) acetylated Histone 3 (**D**), Histone 3 K36-trimethyl (**E**) and Histone 3 K27-trimethyl (**F**) enrichment at the HIV-1 promoter in control (shControl) and PCID2-knockdown (shPCID2) J-Lat 11.1. Total H3 at the HIV-1 LTR is represented as % input. H3Ac, H3K36me3 and H3K27me3 enrichment at the HIV-1 LTR is represented as relative enrichment normalized to Total H3 as shown in A and relative to control. Bars and error lines represent, respectively, mean and SEM of four ChIP-qPCR experiments in four independent chromatin preparations and shRNA-mediated knockdown experiments except for H3K36me3 mark where ChIP-qPCR was performed three times.

We next examined the role of PCID2 as a repressor of HIV-1 transcription during latency by probing the enrichment or loss of, respectively, activating and repressing histone marks at the HIV-1 promoter in presence or absence of PCID2. We performed ChIP-qPCR for total Histone 3, Histone 3 Acetyl (H3Ac) and H3K36 trimethyl (H3K36me3) as a marker for active transcription and H3K27 trimethyl (H3K27me3) as a marker of transcriptional repression and analyzed their presence at the LTR in J-Lat cells expressing or depleted of PCID2. Our data shows that, when PCID2 is depleted, there is an enrichment of H3Ac and H3K36me3 at the HIV-1 promoter (Figure 2C-E) indicating a release in transcriptional blocks and consistent with an active transcriptional state ^29,30^. Concomitant with this, we also observed a decrease in the repressive H3K27 trimethyl histone mark at the HIV-1 promoter in PCID2 knockdown cells (Figure 2F) ^30^. Altogether, our data shows that reduced PCID2 levels lead to a re-organization of the HIV-1 LTR chromatin landscape consistent with increased transcriptional activation, confirming that PCID2 maintains HIV-1 latency by acting on blocks at transcription initiation.

### PCID2 represses HIV-1 gene expression at post-transcriptional steps of gene regulation and interacts with proteins involved in multiple steps of RNA processing

Once transcription is initiated, nascent RNA is co-transcriptionally packed into messenger ribonucleoparticles (mRNPs) for its efficient trafficking and export. The TREX-2 complex then facilitates transport of export-competent mRNPs to the nuclear pore complex ^20–22^. Surveillance, trafficking and export of HIV-1 viral RNA containing mRNPs are tightly regulated during viral latency ^4^. It has recently been shown that the main blocks to HIV-1 gene expression in cells obtained from PLWH occur at post-transcriptional steps ^11,12^. Hence, in order to assess the possible contribution of PCID2 in promoting HIV-1 latency at a post-transcriptional level we analyzed the release of blocks at this specific step in PCID2 depleted J-Lat 11.1 cells with a transcriptional profiling assay as described in Figure 3A. For this, we calculated the relative change in the ratios of Tat-Rev RNA (completely spliced viral RNA) over U5 UTR region. Upon PCID2 knockdown we observe a significant increase in Tat-Rev/UTR RNA ratios (Figure 3B), indicative of a release in blocks at post-transcriptional steps of gene expression.

**Figure 3.**
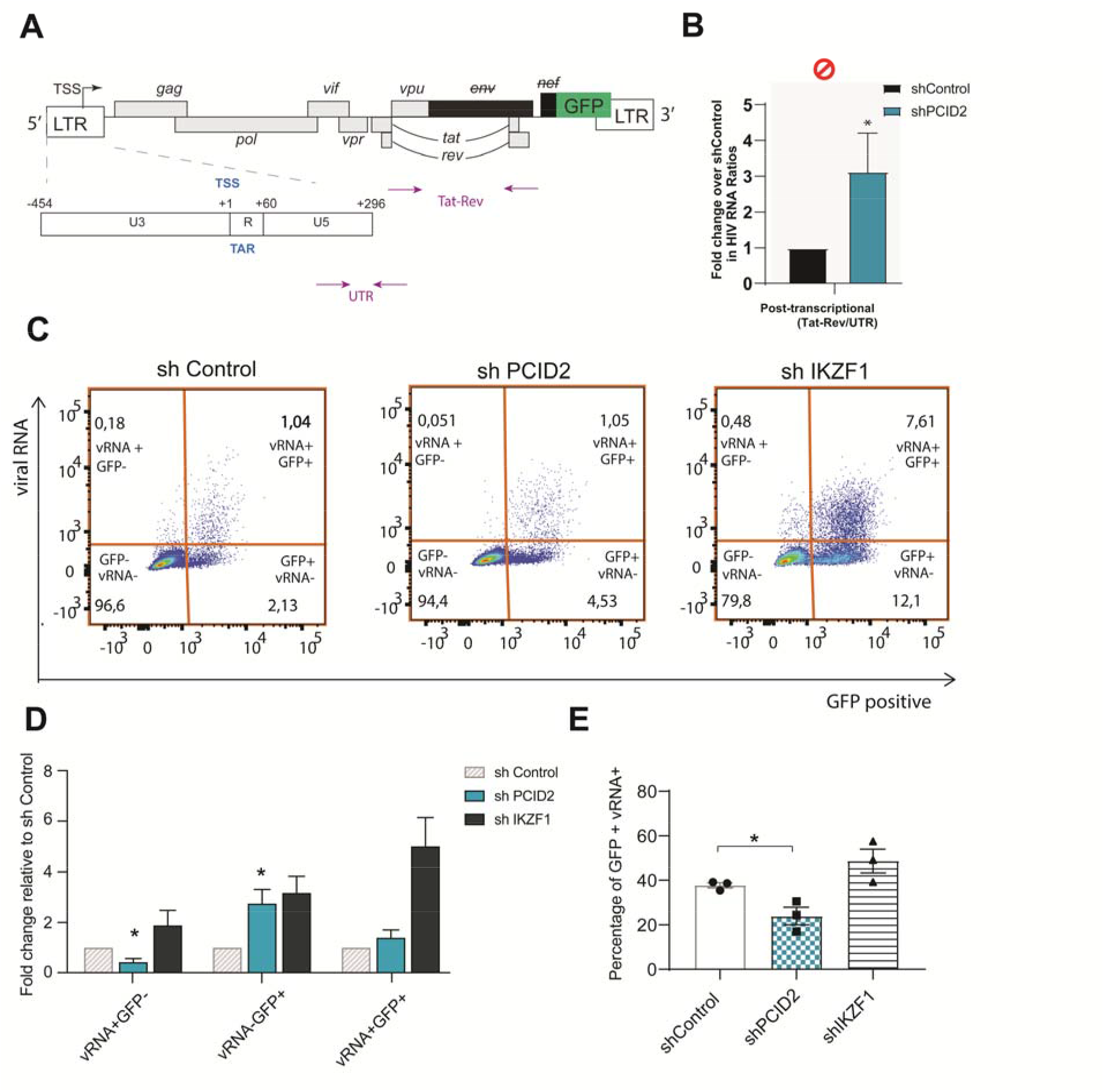
PCID2 blocks post-transcriptional steps of HIV-1 latency. **A**. Schematic figure of the HIV-1 genome in J-Lat 11.1. Arrows represent primer location and amplicons across the HIV-1 genome used in the transcriptional profiling assay. **B**. Transcriptional profiling assay of shControl and shPCID2 J-Lat 11.1. Gene expression blocks at post-transcriptional steps are assessed by calculating the ratio of the relative abundance of HIV-1 RNA species as shown figure and A. Data is presented as fold change in HIV-1 RNA ratios in shPCID2 relative to shControl. Bars represent mean of 5 independent shRNA-mediated knockdown experiments and error lines represent SEM. Statistical significance was determined by t-test; *p<0.01. **C**. Representative FISH-Flow plots of shControl, shPCID2 or shIKZF1 knockdown J-Lat 11.1 cells. **D**. Bar graph showing fold change in vRNA+GFP-, vRNA-GFP+ and vRNA+GFP+ percentages in shControl, shPCID2 and shIKZF1 cells. Bars and error lines represent mean and SEM (n=3). Statistical significance was determined by t-test, * p<0.05. E. Percentage of GFP+ cells that express viral RNA (vRNA+) in shControl, shPCID2 and shIKZF1 as analysed by FISH-Flow. Statistical significance was calculated by t-test; *p<0.05.

To better distinguish between the role of PCID2 in transcription and post-transcriptional steps of HIV-1 gene expression regulation, we performed FISH-Flow, a technology that discriminates between and quantifies transcriptional and post-transcriptional proviral activity at a single cell level, in shControl and shPCID2 J-Lat 11.1 cells (Figure 3C-E, Supplementary Figure 4). FISH-Flow uses probes specific for the GagPol region to measure unspliced HIV-1 viral RNA (US vRNA) to characterize LTR transcriptional activity by flow cytometry. Simultaneously, GFP, that replaces the nef gene in J-Lat cells, is a translation product of completely spliced HIV-1 viral RNA (CS vRNA) and can be measured as a marker of early viral reactivation and viral RNA splicing. In latent cells, and consistent with literature, there is a basal level of residual transcription and basal expression of GFP ^31^ (Figure 3C-D), with about 37.7% (mean) of GFP+ cells also expressing vRNA (Figure 3E). To demonstrate the effect of a release in a transcriptional block specifically, we performed FISH-Flow in J-Lat 11.1 cells shRNA depleted of Ikaros zinc finger 1 (IKZF1), a recently identified HIV-1 transcriptional repressor ^16^. Indeed, we observed strong viral reactivation in IKZF1-knockdown cells as shown by a marked increase in vRNA and GFP producing cells (Figure 3D), of which 48.6% (mean) also express vRNA (Figure 3E). Interestingly, when PCID2 is depleted, we observed a significant decrease of vRNA+ GFP-cells and an increase in cells only expressing GFP (Figure 3D). In addition, the percentage of GFP+ cells that also express vRNA was significantly reduced (23.8%, mean) (Figure 3E). This indicates that, while depletion of PCID2 in J-Lat cells leads to early viral reactivation, there is also an accompanying misregulation in splicing, resulting in a decrease in intron-containing viral RNA in both GFP– and GFP+ cell populations, but an overall increase in GFP+ cells, indicative of increased abundance of completely spliced HIV-1 vRNA species.

These results point to a potential role for PCID2 in regulation of HIV-1 latency at distinct post-transcriptional steps of gene expression, such as splicing and export. Although PCID2 and the TREX2 complex have mostly been studied for their role in linking transcription with transport of export-competent mRNPs, it is currently unknown whether other steps of RNA processing are influenced by PCID2. Actually, members of the THO/TREX complex, that interact with TREX-2 and also mediate transcription with mRNP export ^32^, are known to interact with and recruit splicing factors to mRNPs ^33,34^. Hence, in order to identify possible molecular pathways by which PCID2 regulates HIV-1 gene expression at a post-transcriptional level in an unbiased manner, we performed immunoprecipitation of exogenously expressed PCID2-Flag in J-Lats 11.1 cells coupled with semi-quantitative mass spectrometry analysis (IP-MS). As a control we used J-Lat cells transfected with an empty vector. We performed the IP-MS pipeline two times with independent PCID2-Flag immunoprecipitations (Figure 4A, Supplementary Figure 5A, Supplementary Table 1) followed by application of stringent unbiased filtering to decrease the possibility of enrichment of false interactors (Supplementary Figure 5B). The raw hits found in both conditions (Control and PCID2-Flag) were filtered for common contaminants and presence of unique peptides >1. Only the hits present in the PCID2-Flag IP with a minimum 1.2 fold threshold were taken into consideration for further analysis. After filtering, 75 and 202 proteins were found in the two independent PCID2-Flag IP-MS runs and 30 proteins are common to both experiments (Figure 4B). As expected, the top hit found in both PCID2-Flag IP-MS runs was PCID2 (Supplementary Table 1). Importantly, protein LENG8, recently suggested to be a mammalian orthologue for yeast factor Sac3 (also named GANP) and associated with PCID2 and other members of the TREX2 complex in vitro ^35^, was found in both IP-MS runs to be one of the top hits uniquely present in PCID2-FLAG IP samples (Figure 4C, Supplementary Table 1). In the common PCID2-Flag IP-MS interactor list (Figure 4C), PCID2 interactors included those involved in mRNA metabolism pathways such as translation (ribosomal factors such as RPLP0 and RPS26, RPS6), mRNA splicing (spliceosome complex proteins PRPF8 and BUD13), and mRNA surveillance and stability (UPF1). We also found PCID2 interacting proteins involved in DNA repair, protein stability, cell response to stress, signal transduction and transcription.

**Figure 4.**
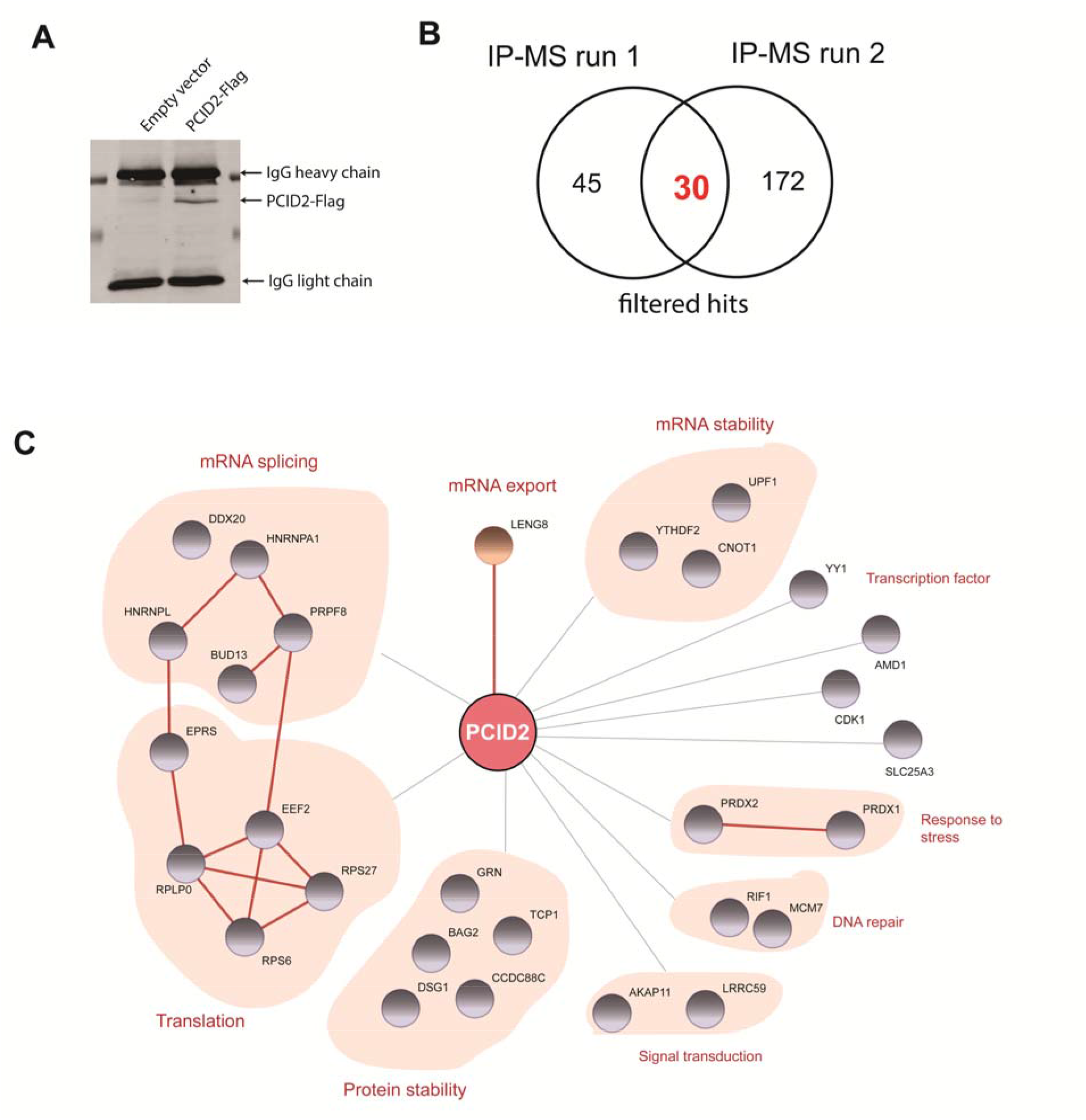
PCID2 interacts with proteins involved in multiple stages of RNA processing. **A**. IP-western blot of PCID2-Flag in control and stably PCID2-Flag expressing J-Lat 11.1 samples for subsequent mass spectrometry analysis. **B**. Venn diagram of the total individual and common number of identified hits by mass-spectrometry in PCID2-Flag over control samples from two independent IP-MS runs. **C**. Network of PCID2 interacting hits identified by mass spectrometry. Hits are classified by cellular function. Grey lines represent PCID2 interacting proteins as analyzed by IP-MS in this study. Red lines represent protein-protein interactions predicted by STRING analysis on experimental evidence only.

Thus, in addition to its role in transcriptional repression of HIV-1 gene expression, our data suggested a role for PCID2 in maintenance of HIV-1 latency at post-transcriptional steps of HIV-1 gene expression via interaction with proteins involved in RNA processing pathways including RNA export, splicing, surveillance and translation.

### PCID2 inhibits alternative splicing during HIV-1 latency

In line with our previous data, the list of PCID2 interactors discovered in our IP-MS approach suggested that, besides RNA export, PCID2 takes part in other steps of RNA trafficking and processing such as splicing. In support of this we found, amongst the identified PCID2 interactors, several proteins involved in RNA splicing, including members of the spliceosome complex (PRPF8, DDX20) and HNRNP family members, known to act as negative regulators of HIV-1 vRNA splicing (Figure 4C) ^36–38^. To determine if PCID2 indeed plays a role in HIV-1 alternative splicing, we performed a PCR-based splicing assay as previously described ^39^ to measure the relative abundance of the three main HIV-1 viral RNA splicing variants: full length US, single spliced (SS) and CS HIV-1 vRNA species. HIV-1 splicing is tightly regulated during latency and US HIV-1 RNA species are more abundant than spliced HIV-1 vRNA species (reviewed in ^40^). Upon PCID2 knockdown, we observed a relative decrease in unspliced vRNA accompanied by a significant increase in completely spliced HIV-1 vRNA (Figure 5A) and a shift in the abundances of unspliced/spliced viral RNA (Figure 5B), in agreement with the data we observed by FISH-Flow (Figure 3C-E). Overexpression of PCID2 in Jurkat lines transfected with an env deficient full-length HIV-1 plasmid shows the opposite effect, with a significant increase in US vRNA and decrease in CS vRNA species (Figure 5C).

**Figure 5.**
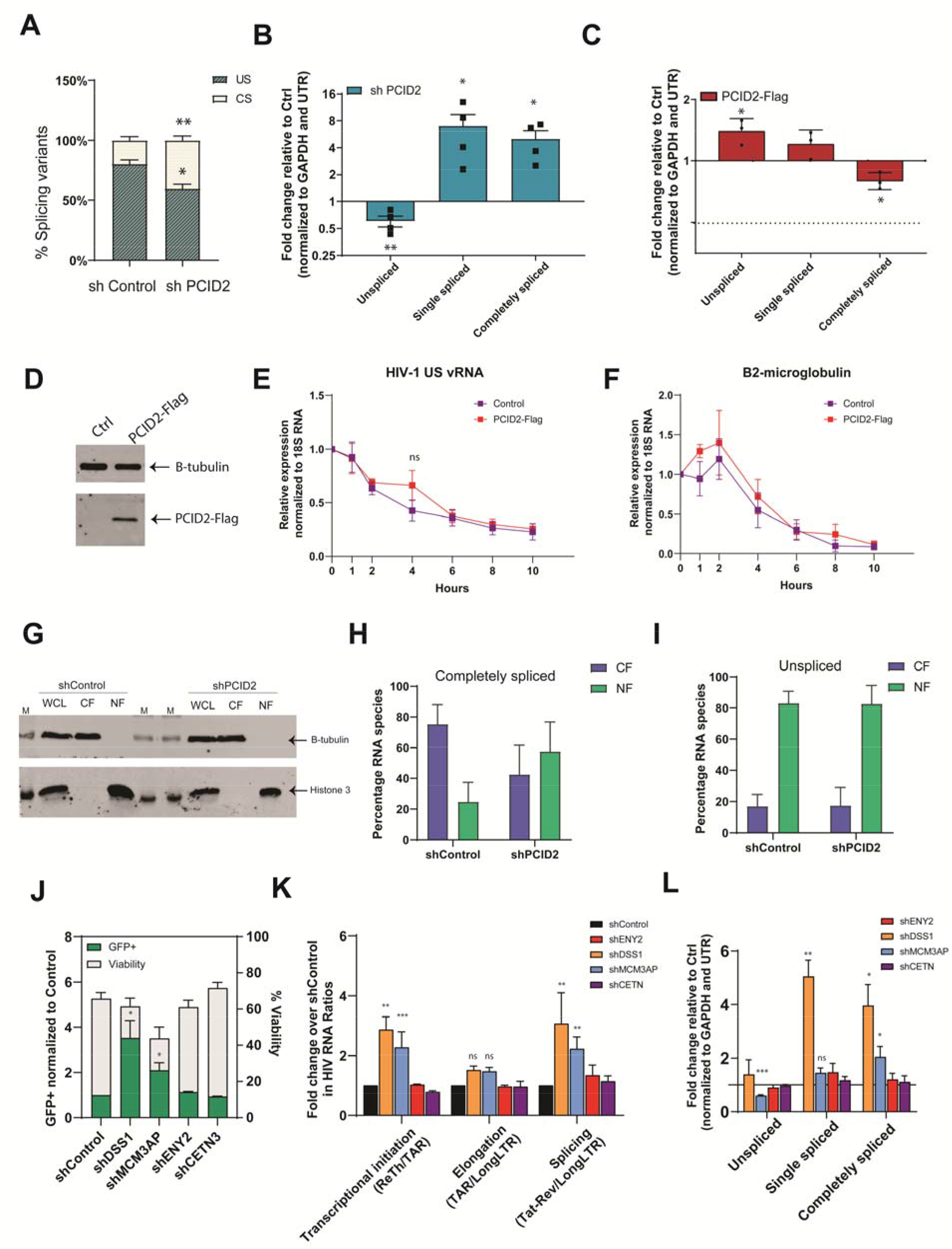
Effect of PCID2 and other TREX2 complex members on transcriptional and post-transcriptional steps of HIV-1 gene expression during viral latency. **A**. Relative percentage of intron-containing unspliced (US) and completely spliced (CS) HIV-1 RNA in shControl and shPCID2 knockdown J-Lat 11.1 **B-C**. Fold change in HIV-1 RNA splicing variants upon PCID2 knockdown relative to shControl (**B**) and in transiently co-nucleofected Jurkat cells with pNL4.3.Luc.R-E-, pRL Renilla and either pBud-Control or pBud-PCID2-Flag (**C**). Values were normalized to GAPDH and UTR HIV-1 transcript. Bars represent mean and error lines represent SEM. Statistical significance was determined by t-test; *p<0.05, **p<0.01. **D**. Western blot of PCID2-Flag in control or PCID2-Flag overexpressing J-Lat 11.1. Beta tubulin was used as loading control. E. HIV-1 unspliced (US) RNA decay dynamics in control and PCID2-Flag overexpressing J-Lat 11.1. Cells were treated with Actinomycin D (10 µg/mL) and RNA was collected at 0, 1, 2, 4, 6, 8 and 10 hours after treatment. Gene expression of unspliced RNA was normalized to 18S RNA and values are calculated relative to timepoint 0 hours. Symbols represent the mean of three independent experiments and error bars represent SEM. Statistical significance was determined by t-test; ns= not significant. **F**. B2-microglobulin RNA decay dynamics in control and PCID2-Flag overexpressing J-Lat 11.1 as described in B. Gene expression of B2-microglobulin RNA was normalized to 18S RNA and values are calculated relative to timepoint 0 hours. Symbols represent the mean of three independent experiments and error bars represent SEM. **G**. Western blot of whole cell lysate (WCL) and cell fractions corresponding to cytoplasm fraction (CF) and nuclear fraction (NF) in shControl or shPCID2 knockdown J-Lat 11.1 to probe for B-tubulin and Histone 3. M refers to protein marker. **H-I**. Percentage of RNA species unspliced HIV-1 vRNA (**H**) and completely spliced HIV-1 vRNA (I) in the cytosolic fraction (CF) and nuclear fraction (NF). Bars and error lines represent mean and SEM (n=3). J. Fold increase in the % of GFP (left y axes) and viability (right y axes) in shControl or shDSS1, shMCM3AP, shENY2 and shCETN3 J-Lat 11.1 cells as measured by flow cytometry. Bars represent mean of three independent shRNA-mediated knockdown experiments and error lines represent SEM. Statistical significance was determined by t-test; *p<0.05. **K**. Transcriptional profiling assay of shControl and shDSS1, shMCM3AP, shENY2 and shCETN3 J-Lat 11.1 cells. Gene expression blocks at transcriptional initiation, elongation, and post-transcriptional steps are asessed by calculating the ratio of the relative abundance of HIV-1 RNA species as shown in the figure. Data is presented as fold change in HIV-1 RNA ratios in knocked down cells relative to shControl. Bars represent mean of three to five independent shRNA-mediated knockdown experiments and error lines represent SEM. Statistical significance was determined by ANOVA test; *p<0.05, **p<0.01, ***p<0.001. **K**. Fold change in HIV-1 RNA splicing variants upon DSS1, MCM3AP, ENY2 and CETN3 knockdown relative to shControl. Values were normalized to GAPDH and UTR HIV-1 transcript. Bars represent mean and error lines represent SEM. Statistical significance was determined by t-test; *p<0.05, **p<0.01, ***p<0.001.

Arguably, the effect on abundance of the different vRNA splicing variants upon PCID2 depletion could also result from PCID2-mediated stabilization of US HIV-1 vRNA. We found, in fact, several proteins that interact with PCID2 involved in RNA surveillance and stability, such as UPF1, previously identified to regulate HIV-1 latency at a post-transcriptional level ^31^. Hence, we next aimed to assess whether the PCID2-mediated change in the relative abundance of splicing variants is explained by a role of PCID2 in HIV-1 RNA stability and decay during latency. To test this, we inhibited general transcription in PCID2-Flag overexpressing and control J-Lat 11.1 lines (Figure 5D-F) by treatment with the DNA intercalator Actinomycin D and measured the relative decay of US HIV-1 vRNA over time. We observed a comparable steady decay both in PCID2-Flag overexpressing and control J-Lat 11.1 lines (Figure 5E). As control for mRNA decay, we also measured decay of B2-microglobulin mRNA and observed, similar to that for US HIV-1 vRNA, a steady decay both in PCID2-Flag overexpressing and control J-Lat 11.1 lines (Figure 5F). These data show that PCID2 has no apparent role in US HIV-1 vRNA stability and that the effect we observe on the abundance and ratios of HIV-1 vRNA splicing variants upon PCID2 knockdown or overexpression result from its role as an inhibitor of viral RNA splicing.

Taken together, our results indicate that PCID2 promotes HIV-1 latency at multiple gene expression regulatory steps; transcriptionally, PCID2 blocks HIV-1 transcription initiation, and post-transcriptionally, PCID2 inhibits alternative splicing.

### TREX2 complex subunits MCM3AP and DSS1 block transcription and misregulate splicing resulting in repression of HIV-1 gene expression

While transcriptional control by PCID2 has been studied as an independent subunit, its involvement in regulation of RNA processing has been almost exclusively linked to its activity as part of the TREX2 complex. Mammalian TREX2 complex is formed by the PCI-containing subunits MCM3AP, DSS1 and PCID2 that form the scaffold and RNA binding platform of the TREX2 complex and subunits ENY2 and CETN2/3,that form the nuclear pore binding dock. TREX2 is hence recruited to nascent RNA and facilitates export of mRNPs via the NXF1 pathway^19–22,41^. We first addressed to what extent viral RNA export is regulated by PCID2 and examined the distribution of intron-containing and completely spliced HIV-1 vRNA species in the nuclear and cytoplasmic compartments upon PCID2 depletion. Indeed, knockdown of PCID2 causes partial nuclear retention of completely spliced HIV-1 vRNA species, as observed by a shift in the distribution between the nuclear and cytoplasmic fractions (Figure 5G-H). This is in line with the role of the PCID2/TREX2 complex in mRNA export, as only completely spliced HIV-1 vRNA variants, because of their small size, are exported via the NXF1 pathway, while intron-containing vRNAs need the viral protein Rev (encoded by CS HIV-1 vRNA) for efficient export ^4^. Accordingly, upon PCID2 knockdown the relative distribution of US HIV-1 vRNA between the cytosolic and nuclear fraction remains stable (Figure 5H), as US HIV-1 vRNA is retained in the nucleus during viral latency (Figure 5I).

Still, the question remains whether the observed suppression of HIV-1 proviral transcription and/or viral RNA alternative splicing by PCID2 occur independently or within the platform of the TREX2 complex. In fact, we previously performed an insertional mutagenesis genetic screen in latent cells and identified the TREX2 complex member MCM3AP as a host factor involved in maintenance of HIV-1 latency ^15^. Hence, to determine the potential role of the TREX2 complex as a whole in suppressing HIV-1 gene expression, we assessed the overall contribution of the other TREX2 complex members to HIV-1 latency. We therefore depleted the individual subunits via shRNA knock-down in J-Lat 11.1 cells and interrogated latency reversal at multiple levels of HIV-1 gene expression (Figure 5J-L, Supplementary Figure 6). We found that depletion of TREX2 nuclear pore binding subunits ENY2 and CETN3 did not lead to HIV-1 reactivation from latency in J-Lat 11.1 (Figure 5J). On the contrary, significant reactivation of HIV-1 latency, as assessed by production of GFP both at the protein and RNA level, was observed consequent to depletion of endogenous DSS1 and MCM3AP that, together with PCID2, form the TREX2 RNA binding dock (Figure 5J).

When looking at the relative release of specific blocks to HIV-1 gene expression by a PCR-based transcriptional profiling assay, we observe that depletion of endogenous DSS1 and MCM3AP, but not ENY2 or CETN3, led to release of blocks in transcription initiation and alternative splicing, similar to what we observed upon PCID2 depletion (Figure 5K). The effect of DSS1 and MCM3AP on suppressing splicing is also reflected by distinct changes in the abundances of viral RNA splicing variants with significant increase in spliced viral RNA upon depletion, while, consistent with our previous data, depletion of ENY2 and CETN3 does not lead to changes in the abundance of viral RNA splicing variants in J-Lat 11.1 (Figure 5L).

Our results are consistent with the notion that PCID2 function in NXF1-mediated mRNA export, and in regulation of HIV-1 latency at transcriptional and post-transcriptional levels of HIV-1 gene expression occur in context of and within functional PCID2-DSS1-MCM3AP RNA-binding docking sub-unit of the TREX2 complex.

## Discussion

In a previous study, we discovered novel LTR-bound regulators of HIV-1 transcription by means of locus specific chromatin immunoprecipitation coupled to mass spectrometry (Catchet-MS) in latent and active infected cells ^16^. Here, we delineate the functions of PCID2, a novel factor we identified to be differentially bound to the latent HIV-1 promoter, as a multifactorial protein that blocks distinct transcriptional and post-transcriptional steps of HIV-1 gene expression to promote viral latency.

In latent cells, endogenous PCID2 is bound to the HIV-1 LTR and removed upon transcriptional activation triggered by PMA treatment, consistent with our findings by the Catchet MS pipeline ^16^. We demonstrate that PCID2 acts as a repressor of HIV-1 transcription initiation during latency and prevents latency reactivation (Figure 1). In literature, PCID2 has been shown to regulate gene expression at multiple levels. The main role of PCID2 described in literature concerns co-transcriptional mRNA processing, and is linked to its function as part of the Transcription and Export Complex 2 (TREX2), that couples transcription with transport of mRNPs containing nascent RNA to the nuclear pore complex for its subsequent export to the cytoplasm ^19–23^. Independent of its role as part of the TREX2 complex, PCID2 has been shown to have a role in genome and protein stability ^24,25^, regulation of embryonic stem cell development and transcriptional regulation of specific genes in hematopoietic stem cells ^27,28^. At a transcriptional level, Ye et al. showed that PCID2 interacts with ZNHIT1, a member of the SRCAP complex, and silences transcription of genes involved in lymphoid fate commitment ^28^. This study also demonstrated that absence of PCID2 increases chromatin accessibility at promoters of lymphoid commitment genes. In agreement, we find that PCID2 promotes latency by enforcing blocks at transcription initiation specifically (Figure 2). Downregulation of endogenous PCID2 re-starts LTR-driven transcription in latent cells and results in changes in the epigenetic state of the promoter as shown by enrichment and decrease in enrichment of, respectively, activating H3Ac and H3K36me3 and repressive H3K27me3 histone marks at the LTR.

The yeast homologue TREX2 complex has been characterized in literature to affect RNA-Pol II mediated transcriptional elongation and absence of several members, including the PCID2 orthologue Thp1, results in significant decrease in elongation of nascent RNA ^42,43^. Transcription elongation is tightly regulated during HIV-1 latency where, in absence of viral protein Tat, RNA-Pol II pauses a few nucleotides after the transcription start site as it encounters the secondary RNA element TAR, and elongation of nascent RNA transcripts is prohibited ^5,44^. Interestingly, in our system, PCID2 depletion does not lead to an increase in elongated viral RNA transcripts. The functions of the mammalian PCID2, however, may very well differ from its yeast orthologue, at least in context of transcriptional elongation at the HIV-1 promoter. Future studies are needed to address the overall role of PCID2 independently or as part of the TREX2 complex on RNA-Pol II mediated transcription elongation in metazoans.

In addition to silencing HIV-1 transcription initiation, our findings demonstrate that PCID2 enforces blocks at post-transcriptional steps of latency (Figure 3-6). Co– and post-transcriptional regulation of HIV-1 gene expression are crucial steps during the establishment and maintenance of viral latency and include blocks at 5’capping, alternative splicing, nucleocytoplasmic export, vRNA surveillance, translation and assembly ^4^. We distinctively show that knockdown of PCID2 in latently infected cell lines mis-regulates the abundance of intron-containing and CS vRNA species, resulting in a marked decrease in levels of US vRNA and accumulation of spliced vRNAs. Our data also demonstrates that this effect is not a consequence of a decrease in stability of US vRNA upon PCID2 depletion, but rather is the result of a release in blocks at alternative splicing. This is further supported by our finding that PCID2 interacts with several members of the spliceosome complex and other splicing modulators such as PRPF8, HNRNPA1 and BUD13. A similar over-splicing effect has been reported in literature as a result of altering the abundance or functionality of SR protein kinases ^45,46^, and absence or mutation of cis-acting elements that prevent binding of the spliceosome complex (reviewed in^47^).

**Figure 6.**
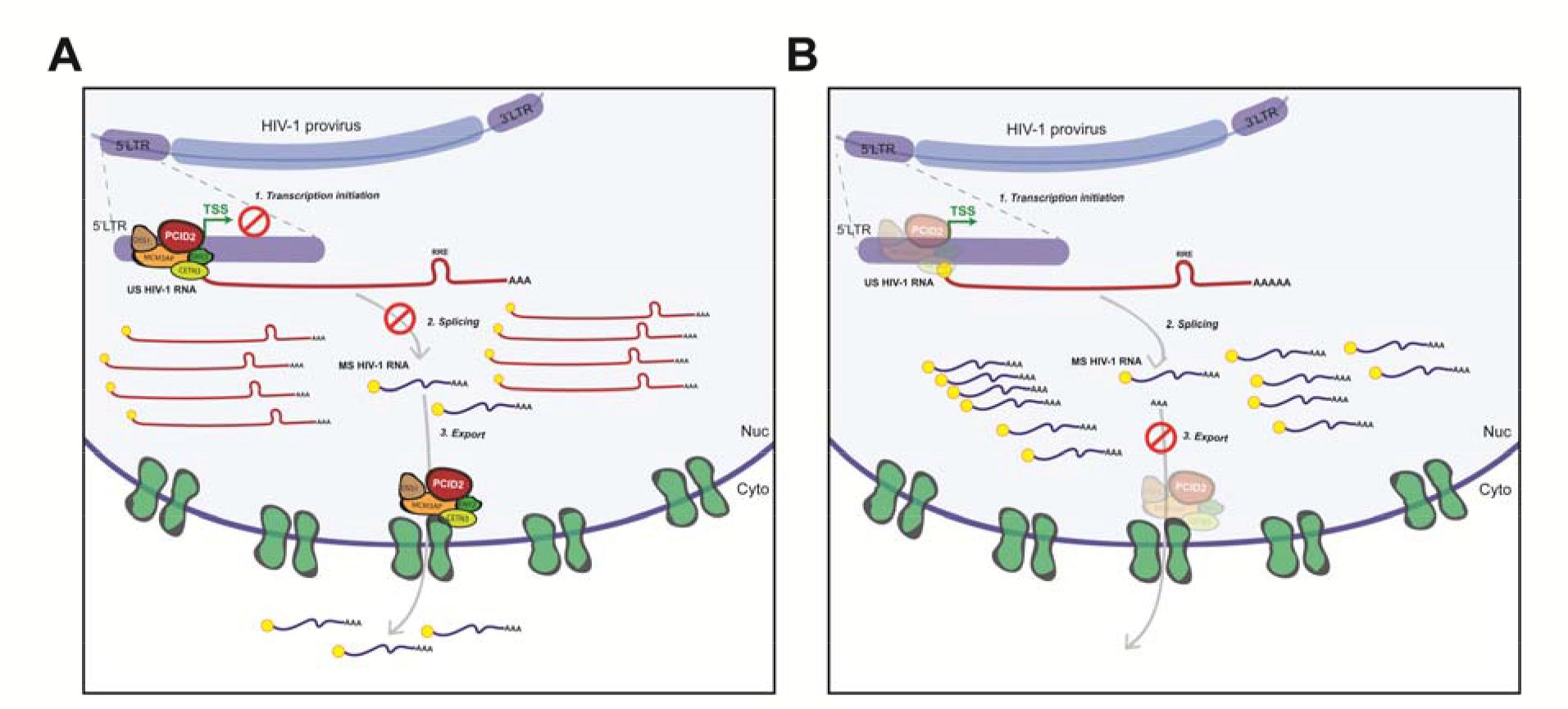
Proposed model for the role of PCID2/TREX2 in HIV-1 gene expression during viral latency. **A**. In latent cells, PCID2/TREX2 is present at the HIV-1 LTR transcription start site (TSS) and blocks HIV-1 gene expression and promotes viral latency at two distinct levels: transcription initiation and RNA alternative splicing. **B**. Upon PCID2-DSS1-MCM3AP downregulation, LTR-driven transcription is re-started. In addition, absence of PCID2-DSS1-MCM3AP leads to increased splicing of intron-containing unspliced (US) viral RNA and accumulation of completely spliced (CS) HIV-1 RNA. Because PCID2 facilitates export of mRNPs as part of the TREX2 complex via the NXF1 pathway, its downregulation leads to nuclear retention of CS HIV-1 vRNA.

The role of PCID2 in RNA metabolism has been primarily linked to its function in nucleocytoplasmic RNA export and other putative functions of PCID2 alone or in context of the TREX2 complex in prior or further steps of RNA processing have been unexplored. Notably, another TREX2 complex member, protein Sem1 (yeast homologue for DSS1) has been shown previously to influence mRNA alternative splicing in yeast ^48^. This raised the question of whether PCID2 exerts its function in transcriptional and/or post-transcriptional steps of HIV-1 gene expression regulation independent or as part of the TREX2 complex. Interestingly, when we examined the relative contribution of the other TREX2 complex members to HIV-1 latency individually, we observed a prominent role for TREX2 subunits MCM3AP and DSS1 in regulating viral latency. This agrees with our previous study in which, by performing an insertional mutagenesis genetic screen in latent cells, we identified MCM3AP as a putative host factor that enforces HIV-1 latency ^15^. We further show that, mirroring the effects observed with PCID2, suppression of HIV-1 gene expression by MCM3AP and DSS1 results from hindered transcription initiation and inhibition of viral RNA alternative splicing. In yeast, Sac3 (yeast homologue for MCM3AP) and Sem1 (DSS1) are also PCI-containing proteins that bind Thp1 (PCID2), and respectively, form the scaffold and stabilize Thp1 (PCID2) in the Sac3-Thp1-Sem1 complex ^49,50^. The sub-complex DSS1-MCM3AP-PCID2 simultaneously form part of the RNA binding platform of the larger TREX2 complex. Our findings imply a specific role for the PCID2-DSS1-MCM3AP sub-complex in interacting with viral RNA and modulating viral latency at both transcriptional and post-transcriptional steps of gene expression ^20,22,49^. In our experimental system we did not observe a clear effect on HIV-1 latency reactivation upon depletion of TREX2 subunits ENY2 and CETN3 that bind the nuclear pore complex for efficient nucleocytoplasmic export of mRNP complexes ^20–22^. Thus, these observations are consistent with the hypothesis that repression of HIV-1 gene expression by PCID2 is an RNA mediated process that likely occurs in context of the PCID2-DSS1-MCM3AP sub-complex of the TREX2 complex and is dependent on nucleic acid binding.

Lastly, a well-described role for PCID2 is its role as part of the TREX2 complex in nucleocytoplasmic export of spliced mRNAs via the NXF1-NXT1 pathway ^20,22,23^. During HIV-1 infection, only CS vRNAs, because of their 2kb size, are exported via NXF1-NXT1. Larger, intron-containing vRNA species need viral protein Rev and other co-factors for their efficient export to the cytoplasm ^4,51^. In yeast, absence or loss of function of Thp1 (PCID2 orthologue) leads to overall inhibition of spliced mRNA export ^23^. Consistent with this, we show that downregulation of PCID2 causes nuclear retention of CS vRNA but not US vRNA species. Of note, we still observe an increase in GFP, encoded by the CS vRNA species, when PCID2 is depleted from J-Lat cells. This could be explained by the fact that our shRNA-mediated knockdown is not 100% efficient and some residual PCID2 still remains, and/or that other compensatory pathways are in place that sustain export of CS vRNA species to the cytoplasm.

In conclusion, our data supports a model in which PCID2-containing complexes maintain HIV-1 latency at two essential steps of HIV-1 gene expression regulation (Figure 6). In our system, PCID2 acts as a repressor of HIV-1 LTR-driven transcription initiation and blocks viral RNA alternative splicing. Hence, upon PCID2 depletion, transcription initiation from the HIV-1 LTR is re-started and blocks at splicing are released, resulting in higher levels of spliced vRNAs and overall viral reactivation from latency. Finally, our study furthers current knowledge on the molecular pathways involved in HIV-1 latency regulation, emphasizing the role of novel multifactorial host cell factors in governing viral reactivation across various steps, including post-transcriptionally.

## Author contributions

Conceptualization: RC, EN, RJP, SR, TM; Experimental design, formal analysis, methodology and validation: RC, EN, JR, JM, CL, SJ, AG, SK, TWK, WD, DD, JD; Writing – original draft and review and editing: RC, RJP, SR, TM.

## Methods

### Resource availability

#### Lead contact

Further information and requests for resources and reagents should be directed to and will be fulfilled by the lead contact, Tokameh Mahmoudi

#### Materials availability

Plasmids generated in this study are available upon request to lead contact, Tokameh Mahmoudi.

#### Data and code availability

- Mass spectrometry data is available in PRIDE repository and can be accessed using the unique identifier PXD043334.
- This paper does not report original code.
- Any additional information required to reanalyze the data reported in this work paper is available from the lead contact upon request.

### Experimental model details

Jurkat cells and latent HIV-1 infected Jurkat clones J-Lat 11.1, 10.6 and A2 ^52^ were cultured in RPMI-1640 media supplemented with heat inactivated 7% Fetal Bovine Serum and 100 µg/mL Penicillin-Streptomycin at 37°C in a humidified 95% air 5% CO_2_ incubator. Newly generated stable control or PCID2-Flag expressing J-Lat 11.1 cells were cultured in RPMI-1640 media supplemented with heat inactivated 7% Fetal Bovine Serum (FBS) and 100 µg/mL Penicillin-Streptomycin at 37°C in a humidified 95% air 5% CO_2_ incubator. HEK 293T cells were cultured in Dulbecco’s Modified Eagle’s Medium (DMEM) supplemented with heat inactivated 7% Fetal Bovine Serum and 100 µg/mL Penicillin-Streptomycin at 37°C in a humidified 95% air 5% CO_2_ incubator.

### Method details

#### Reagents

Cells were treated as indicated in the figures or figure legends with the following compounds: PMA (phorbol 12-myristate 13-acetate, Sigma Aldrich), Valproic acid (Sigma Aldrich), Actinomycin D (Sigma Aldrich). All chemicals were reconstituted following manufacturer’s instructions.

#### Western blot

1-2 million cells (up to 5 million cells for PCID2 knockdown western blot) were lysed using an NP-40 IP lysis buffer (1% NP-40, 25mM Tris pH 7.4, 150mM NaCl, 1mM EDTA, 5% glycerol, 1 U/mL EDTA-free protease inhibitor cocktail (Roche) and 1mM DTT) for 30 minutes on ice and centrifuged for 10 minutes at 14000 rpm in a cold centrifuge. Supernatants were collected, 1x Laemmli loading buffer was added and lysates were boiled for 5 min at 95°C and subjected to 12% SDS-PAGE separation. The following antibodies were used for detection of proteins by western blot: anti-M2-FLAG (Sigma, F3165), anti-β-tubulin (Sigma, T5168), anti-PCID2 (Genetex, GTX52023) and anti-Histone 3 (Abcam, ab1791) antibodies.

#### Lentiviral shRNA-mediated knockdown

Lentiviral constructs containing the shRNA targeting the gene of interest were obtained from the MISSIONx shRNA library (Sigma), facilitated by Erasmus Center for Biomics (Key Resource Table). Plasmids were amplified in DH5α bacterial cells and isolated with Invitrogen™ PureLink™ HiPure Plasmid Miniprep Kit. Pseudotyped lentivirus were obtained by co-transfecting LJ 7×10^6^ HEK293T cells with 6 µg of lentiviral construct with 4,5 µg of envelope plasmid pCMVΔR8.9 and 2 µg of packaging plasmid pCMV-VSVg mixed with 10 mM polyethyleneimine containing serum free-DMEM (transfection mix was incubated 15 min at room temperature) for 12 hours. Medium was then replaced with RPMI-1640 supplemented with 7% Fetal Bovine Serum (FBS) and 100 µg/mL Penicillin-Streptomycin, and supernatant was harvested 36, 48 and 60 hours post-transfection, filtered through a cellulose acetate membrane (0.45 µm pore) and stored at –80°C for later use.

#### Establishment of stable PCID2-Flag expressing lines

Stable PCID2-Flag expressing J-Lat 11.1 cell lines were generated by cloning the PCID2 open reading frame (ORF) in a pBud backbone (pBud Tag3 C-ter) behind an EF1α promoter and introducing a 2x FLAG tag at the C-term of the PCID2 ORF, using DH5α bacterial cells to propagate the plasmids. The plasmid contains a Bleomycin resistance cassette for bacterial selection and a Geneticin® cassette for mammalian cell selection. Stable cell lines were generated by nucleofecting 2 µg of pBud-PCID2-Flag plasmid or empty pBud as control using Amaxa Nucleofector (Lonza) and Nucleofector Kit R (Lonza) following manufacturer instructions. Briefly, 5×10^6^ were centrifuged at 1500 rpm for 5 min at room temperature and resuspended in 100 µL of solution R, and nucleofected in the presence of the plasmid mix containing pBud control or pBud-PCID2-Flag using program O28. Nucleofected cells were immediately incubated in pre-warmed serum free and antibiotic free RPMI media for 15 to 30 min at 37°C in a humidified 95% air 5% CO_2_ incubator and then transferred to 5 mL of pre-warm RPMI-F7. 4 days after nucleofection cells were selected for two weeks with 0.5 mg/mL Geneticin® and expanded. A clone was selected based on exogenous expression of PCID2-Flag confirmed by western blot and comparable levels of PCID2 mRNA assessed by RT-PCR.

#### Effect of transient PCID2 overexpression on HIV-1 gene expression and latency reactivation

To assess the effect of PCID2 overexpression on HIV-1 reactivation from latency and viral RNA stability as described below, J-Lat 11.1 were nucleofected with 2 µg of pBud-PCID2-Flag plasmid or empty pBud as control as described above. After 4 days cells were selected for two weeks with 0.5 mg/mL Geneticin® and used for further experiments.

To determine the effect of PCID2 overexpression on HIV-1 gene expression, Jurkat cells were co-nucleofected with the following mix of plasmid constructs: 300 and 600 ng of empty pBud control or pBud-PCID2-Flag, 800 ng pNL4.3.Luc.R-.E– (NIH Reagents program) and 200 ng pRL Renilla (Promega E2231) as control for nucleofection efficiency. Nucleofection was performed as described above. After 2 days, cells were processed for luciferase readings or RNA extraction as described below.

#### Dual luciferase assay

One million Jurkat cells transiently expressing pBud control or pBud-PCID2-Flag, pNL4.3.Luc.R-.E– and pRL Renilla were collected 48 hours post-nucleofection and processed for luciferase readings using Dual-Glo® Luciferase Assay System (Promega) following manufacturer instructions. Renilla luciferase activity was used as a nucleofection efficiency control for normalization of Firefly luciferase readings.

#### Chromatin preparation

In order to obtain chromatin, 80-100 million cells (control and untreated or PMA-treated PCID2-Flag expressing cells; shControl or shPCID2 cells) were centrifuged and resuspended in PBS supplemented with 1mM CaCl_2_ and 1mM MgCl_2_ (PBS+/+). Cells were crosslinked by addition of Buffer A (0.1M NaCl, 1mM EDTA, 0.5 mM EGTA, 20mM HEPES) with formaldehyde (molecular grade methanol-free 16% formaldehyde, Polysciences Inc.) at a final percentage of 1% for 30 minutes at room temperature in a vertical rotator. The crosslinking reaction was stopped with 150 mM final concentration of Glycine and cells were washed once with PBS+/+. The pellets were washed consecutively once with cold Buffer B (0.25% Triton x-100, 1mM EDTA, 0.5 mM EGTA, 20mM HEPES pH 7.6) and once with cold Buffer C (150 mM NaCl, 1mM EDTA, 0.5 mM EGTA, 20mM HEPES pH 7.6) for 10 min on a vertical rotator at 4°C and centrifuged for 5 min 1500 rpm at 4°C. Pellets were resuspended in 2.5mL of ChIP Incubation buffer (1% Triton x-100, 150 mM NaCl, 1mM EDTA, 0.5 mM EGTA, 20mM HEPES pH 7.6) supplemented with 1% SDS and protease inhibitors (EDTA-free protease inhibitor cocktail; Roche) and transferred to 15 mL polystyrene Falcon tubes compatible with sonication. Chromatin was sonicated with Bioruptor (Diagenode) using 15mL probes for 12-15 cycles of 30 seconds ON 30 seconds OFF intervals at high intensity to achieve fragments in between 100 and 500bp. The sonicated chromatin was centrifuged for 15 min 14000 rpm at 4°C to get rid of cell debris.

To check for chromatin quality and fragment size, 100 µL of chromatin were added to 300 µL of ChIP Elution buffer (1% SDS, 0.1M NaHCO_3_), brought to 200 mM NaCl final concentration, and decrosslinked for a minimum of 4 hours or overnight at 65°C on a heat block with shaker at 750 rpm. DNA extraction was performed using a standard phenol-chloroform isoamyl alcohol isolation protocol. Briefly, an equal volume of phenol:chloroform:isoamyl alcohol was added to the decrosslinked chromatin, mixed vigorously, and centrifuged at 14000 rpm for 5 min at 4°C. The aqueous phase was collected and an equal volume of 24:1 chloroform:isoamyl alcohol was added, mixed thoroughly and spun at 14000 rpm for 5 min at 4°C. The DNA contained in the aqueous phase was then precipitated with 100% ethanol in presence of glycogen as a carrier and NaAc, and snap frozen with liquid N_2_. The DNA pellet was washed once with 70% ethanol, dried, and resuspended in 100 µL nuclease-free DEPC-treated water. Quality and size of the fragments were analyzed on a 1.2% agarose gel.

#### Chromatin immunoprecipitation (ChIP) – qPCR

To assess the enrichment of PCID2 at the HIV-1 LTR in control and exogenously expressing PCID2-Flag cells, we used the equivalent of 20-30 million cells as chromatin input. Chromatin was diluted with ChIP incubation buffer with no SDS to reach a final concentration of 0.15% SDS and pre-cleared overnight with 50 µL of a mix of Protein A/Protein G sepharose beads washed two times with ChIP incubation buffer 0.15% SDS. 100 µL per sample of anti-FLAG^®^ M2 affinity beads (Sigma) were washed twice with ChIP incubation buffer 0.15% SDS and blocked overnight with 0.1% bovine serum albumin and 400 ng/mL of sheared salmon sperm DNA. An aliquot of the pre-cleared chromatin was set aside for quality control and input quantification. PCID2-Flag bound complexes were isolated by precipitating the pre-cleared chromatin with LJ 100µL of blocked affinity beads overnight at 4°C. The day after, beads were washed twice (10 minutes 4°C, 1500 rpm in a vertical rotator) with Buffer 1 (0.1% SDS, 0.1% DOC, 1% Triton x-100, 150mM NaCl, 1 mM EDTA pH 8.0, 0.5 mM EGTA, 20mM HEPES pH8.0) once per buffer with Buffer 2 (500mM NaCl: 0.1% SDS, 0.1% DOC, 1% Triton x-100, 500mM NaCl, 1 mM EDTA pH 8.0, 0.5mM EGTA, 20mM HEPES pH8.0) and Buffer 3 (0.25M LiCL, 0.5%DOC, 0.5% NP-40, 1mM EDTA pH 8.0, 0.5mM EGTA pH 8.0, 20mM HEPES pH 8.0), and twice with Buffer 4 (1mM EDTA pH8.0, 0.5mM EGTA pH 8.0, 20mM HEPES pH 8.0). Immunoprecipitated complexes were then eluted with ChIP Elution buffer (1% SDS, 0.1M NaHCO_3_) for 30 minutes at room temperature in a vertical rotator.

For ChIP of core and modified histones and other factors in untreated and PMA-treated PCID2-Flag expressing cells, or in knockdown lines we used less chromatin input, corresponding to 10-20 million cells. Chromatin was pre-cleared and 100 µL per sample of a mix of Protein A/Protein G sepharose beads were blocked overnight as described above. The day after, bound complexes were isolated by overnight precipitation of the pre-cleared chromatin with 100 µL of blocked beads in combination with 3-4 µg of antibodies (anti-Histone H3 (Abcam ab17913, µg), Histone H3Ac (Millipore 17-615, 3µg), Histone H3K27me3 (Active Motif 39055, 3µg), Histone H3K36me3 (Abcam ab9050, 4µg), washed twice with Buffer 1, once per buffer with Buffer 2 and Buffer 3, and twice with Buffer 4, and eluted with ChIP Elution Buffer as described above.

DNA extraction was performed the day after using a standard phenol-chloroform isoamyl alcohol isolation protocol as described above. Enrichment of proteins, core and modified histones and transcription factors at the HIV-1 LTR was assessed by quantitative PCR using primers spanning the full promoter (Table 1) with GoTaq qPCR Master mix kit (Promega) in a CFX Connect Real-Time PCR thermocycler (BioRad). Relative enrichment over the DNA input was calculated with the 2 ^−ΔCt^ method ^53^.

**Table 1.**
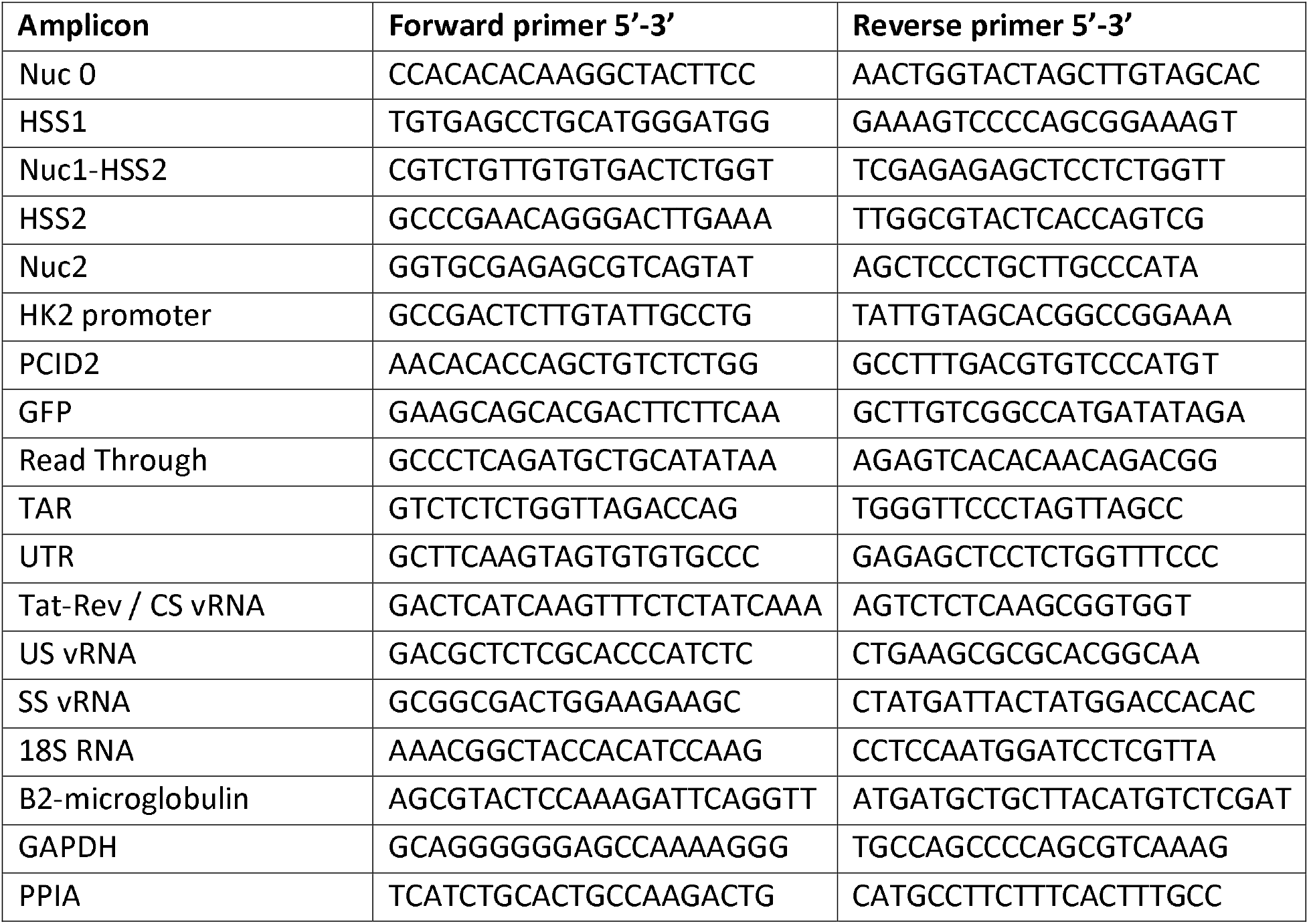
List of primers used for quantitative PCR.

#### PCID2 immunoprecipitation for mass spectrometry

Twenty million control and stable exogenously expressing PCID2-Flag cells were lysed in 1 mL IP lysis buffer (1% NP-40, 25mM Tris pH 7.4, 150mM NaCl, 1mM EDTA, 5% glycerol, 1 U/mL EDTA-free protease inhibitor cocktail (Roche) and 1mM DTT) for 30 min on ice and centrifuged for 10 min 14000rpm at 4°C. Lysates were pre-cleared for 6 hours with 50 µL of a mix of Protein A and Protein G sepharose beads. 80 µL of Anti-FLAG M2 affinity gel per sample were washed twice in IP lysis buffer and incubated overnight with pre-cleared lysates. The day after, beads were washed twice with Wash buffer A (10 mM HEPES, pH 7.4, 10mM KCl, 10mM NaCl, 1 mM MgCl2, 0.05% Nonidet P-40) once with Wash buffer B (10 mM HEPES, pH 7.4, 10mM KCl, 0.07% Nonidet P-40), as described by Lagundzin, D. et al. ^54^, and two times with PBS. Beads were further processed for Mass spectrometry as described below.

#### Mass spectrometry

Proteins were on-bead digested with sequencing grade trypsin (1:100 (w:w), Roche) overnight at room temperature. Protein digests were then desalted using a Sep-Pak tC18 Vac cartridge (Waters) and eluted with 80% acetonitrile (AcN). Peptides were then analyzed by nanoflow LC-MS/MS as described below.

Nanoflow LC-MS/MS was performed on an EASY-nLC system (Thermo) coupled to a Fusion Lumos Tribrid Orbitrap mass spectrometer (Thermo), operating in positive mode and equipped with a nanospray source. Peptide mixtures were trapped on a ReproSil C18 reversed phase column (Dr Maisch GmbH; column dimensions 1.5 cm × 100 µm, packed in-house) at a flow rate of 8 µl/min. Peptide separation was performed on ReproSil C18 reversed phase column (Dr Maisch GmbH; column dimensions 15 cm × 50 µm, packed in-house) using a linear gradient from 0 to 80% B (A = 0.1% FA; B = 80% (v/v) AcN, 0.1 % FA) in 70 or 120 min and at a constant flow rate of 250 nl/min. The column eluent was directly sprayed into the ESI source of the mass spectrometer. All mass spectra were acquired in profile mode. The resolution in MS1 mode was set to 120,000 (AGC: 4E5), the m/z range 350-1400. Fragmentation of precursors was performed in 2 s cycle time data-dependent mode by HCD with a precursor window of 1.6 m/z and a normalized collision energy of 30.0; MS2 spectra were recorded in the orbitrap at 30,000 resolution. Singly charged precursors were excluded from fragmentation and the dynamic exclusion was set to 60 seconds.

Data analysis: Mass spectrometric raw data were analyzed using the MaxQuant software suite (version 2.1.3.0 ^55^ for identification and relative quantification of proteins. A false discovery rate (FDR) of 0.01 for proteins and peptides and a minimum peptide length of 6 amino acids were required. The Andromeda search engine was used to search the MS/MS spectra against the Homo sapiens Uniprot database (version May 2022) concatenated with the reversed versions of all sequences and a contaminant database listing typical background proteins. A maximum of two missed cleavages were allowed. MS/MS spectra were analyzed using MaxQuant’s default settings for Orbitrap and ion trap spectra. The maximum precursor ion charge state used for searching was 7 and the enzyme specificity was set to trypsin. Further modifications were cysteine carbamidomethylation (fixed) as well as methionine oxidation. The minimum number of peptides for positive protein identification was set to 2. The minimum number of razor and unique peptides set to 1. Only unique and razor non-modified, methionine oxidized and protein N-terminal acetylated peptides were used for protein quantitation. The minimal score for modified peptides was set to 40 (default value).

MS raw data and data for protein identification and quantification were submitted as supplementary tables to the ProteomeXchange Consortium via the PRIDE partner repository with the data identifier PXD043334.

The identified hits in both IP-MS runs were first filtered for common contaminants, including filtering of keratin and immunoglobulin proteins. Next, hits were filtered by unique peptides larger than 1. Lastly, a fold enrichment was calculated of iBAQ value in PCID2-Flag samples over control samples, and only hits enriched >1.2 fold were included for further analysis. Common hits for both IP-MS runs were identified and pathway and interaction analysis was performed using STRING database.

#### RNA stability assay

Twenty million control or overexpressing PCID2-Flag lines were plated in 6 mL of RPMI-F7 medium and 10 µg/µL of Actinomycin D was added to the cultures. An aliquot of 1 million cells was collected and lysed in TRI reagent (Sigma) immediately, corresponding to timepoint 0 hours. Cells were then collected and lysed at 1 hour post-treatment and in several sequential timepoints of 2, 4, 6, 8 and 10 hours post-treatment initiation. Total RNA was extracted and 10 µL of RNA was used for cDNA synthesis as described above/below. Decay of HIV-1 unspliced RNA and beta-2-microglobulin was assessed by relative decrease in gene expression from timepoint 0 hours. Values calculated using the 2 ^−ΔΔCt^ and normalized to 18S RNA to account for RNA input variability.

#### FISH-Flow

To analyze the dynamics of viral RNA (vRNA) and/or GFP-producing cells by FISH-Flow, five million control and PCID2-knockdown cells were collected, fixed, permeabilized and processed as described before ^13^ using the PrimeFlow RNA assay (ThermoFisher Scientific) according to manufacturer protocol. Viral RNA producing cells were detected by labelling HIV-1 unspliced mRNA with a set of 40 probe pairs against the GagPol region (Affymetrix eBioscience catalogue number GagPol HIV-1 VF10-10884) and RPL13A (Affymetrix eBioscience, catalogue number VA-13187) as a control diluted 1:5 in diluent provided in the kit and hybridized to the target HIV-1 vRNA for 2 hours at 40°C. As a control for hybridization, we used a probe set against the ribosomal RNA RPP. After hybridization, the excess probes were washed and cells were stored overnight in the presence of RNAsin. The next day, signal was amplified by incubating samples with sequential 1.5 hours, 40°C incubations with pre-amplification and amplification mix, followed by labelling of the amplified RNA with fluorescently-tagged probes for 1 hour at 40°C. Cells were acquired on a BD LSR Fortessa Analyzer and analysis was performed using FlowJo V10 software (Treestar). Gates to quantify viral RNA+ only cells, GFP+ only cells, and vRNA+ GFP+ cells were set using the shControl sample.

#### Confocal imaging

For confocal imaging of viral RNA and GFP in control and PCID2-knockdown cells, we used 100.000 cells processed for FISH-Flow as described above. Cells were dried on a slide, mounted with DAPI-containing Fluorescent Mounting Media (Dako Omnis), and visualized using a LSM700 (Zeiss) confocal microscope.

#### Total RNA isolation and RT quantitative PCR analysis

Cells were lysed with TRI reagent (Sigma) and total cell-associated RNA was isolated following manufacturer instructions. Up to one microgram of RNA was used to synthetized cDNA using High-Capacity cDNA Reverse Transcription Kit (Applied Biosystems 4368814) according to kit protocol. Real time quantitative PCR was performed using GoTaq qPCR Master mix kit (Promega) in a CFX Connect Real-Time PCR thermocycler (BioRad). Normalized relative gene expression was calculated with the 2 ^−ΔΔCt^ method using PPIA as reference gene. Primer list is available in Key Resource Table.

#### Subcellular fractionation

To assess the relative abundance of HIV-1 viral RNA species in the nucleus and cytoplasm of shControl and shPCID2 J-Lat 11.1 cells, we used a modified version of REAP protocol by Suzuki et al ^56^. 2 million cells were lysed in 900 µL of PBS 0.1% NP-40 and mechanically triturated for 4-5 times with a micropipette. 300 µL are taken aside as whole cell lysate of which 100 µL are added to 900 µL of TRI reagent for RNA isolation. The remaining 600 µL are spun for 1 minute at 14000 rpm in a cold centrifuge. 300 µL of the supernatant are set aside as cytoplasmic fraction of which 100 µL are added to 900 µL of TRI reagent for RNA isolation. The rest of the supernatant is discarded and 900 µL of PBS 0.1% NP-40 is added to wash the pellet, which corresponds to the nuclear fraction, for 1 minute at 14000 rpm in a cold centrifuge. Supernatant is discarded and pellet is resuspended in 1 mL of cold TRI reagent. Total RNA was isolated from samples in TRI reagent and processed for cDNA synthesis as described above. RNA concentration of each fraction independently was normalized amongst samples. The relative abundance of unspliced and completely spliced HIV-1 viral RNA species in the nucleus and cytoplasm was calculated with the 2 ^−ΔCt^ method. Ratios were calculated for each vRNA species in the nucleus and cytoplasm and presented as percentages. To check the efficiency of the subcellular fractionation at a protein level, a parallel fractionation protocol was performed as described above except 4x SDS loading buffer with 100 mM DTT is added to the collected whole cell lysate and cytoplasmic fractions. The pellet corresponding to the nuclear fraction is dissolved in 300 µL of 1x SDS loading buffer with 100 mM DTT and sonicated with Bioruptor (Diagenode) 5 cycles of 30 seconds ON 30 seconds OFF intervals at high intensity. Samples were then processed for western blot analysis.

## Quantification and statistical analysis

Statistical analysis was performed as indicated in the figure legends using Graphpad Prism v8.3.0.

## Supplemental Information titles and legends

**Supplementary** Figure 1. **ChIP-qPCR analysis of PCID2-Flag enrichment at the HIV-1 LTR**

**A**. Gene expression analysis of exogenously expressing PCID2-Flag J-Lat 11.1 cells (Stable PCID2-Flag) relative to Control lines (empty vector) and normalized to cyclophilin A. Bars represent mean of 3 separate collections and error lines represent SEM. **B-C**. Enrichment of PCID2-Flag at the HIV-1 LTR expressed as % of input in control and PCID2-Flag expressing J-Lat 11.1 lines as assessed by chromatin immunoprecipitation (ChIP) coupled with qPCR. Primers spanning across the HIV-1 promoter were used to assess relative enrichment of PCID2-Flag at sequential regions of the LTR. HK2 promoter was used as a control genomic region. ChIP-qPCR corresponds to two independent chromatin preparations (**B** and **C**), bars and error lines represent, respectively, mean and SEM of technical duplos. **D**. ChIP-qPCR analysis of total Histone 3 (E) enrichment at the HIV-1 promoter in untreated and PMA treated PCID2-Flag expressing J-Lat 11.1 lines. Total H3 presence at the HIV-1 LTR is represented as % input and is used for normalization of acetylated Histone 3 as presented in Figure 1D. ChIP-qPCR corresponds to one chromatin preparation, bars and error lines represent, respectively, mean and SD of technical duplos. **E-F**. ChIP-qPCR analysis of total Histone 3 (E) and acetylated Histone 3 (F) enrichment at the HIV-1 promoter in untreated and PMA treated PCID2-Flag expressing J-Lat 11.1 lines. Total H3 presence at the HIV-1 LTR is represented as % input. H3Ac enrichment at the HIV-1 LTR is represented as relative enrichment normalized to Total H3 as shown in D. ChIP-qPCR corresponds to one chromatin preparation, bars and error lines represent, respectively, mean and SD of technical duplos.

**Supplementary** Figure 2. **PCID2 knockdown in J-Lat clones 10.6 and A2**

**A**. Fold increase in the % of GFP (left y axes) and viability (right y axes) in shControl or shPCID2 J-Lat 10.6 cells as measured by flow cytometry. Bars represent mean of 3 independent shRNA-mediated knockdown experiments and error lines represent SEM. Statistical significance was determined by t-test; **p<0.01. **B**. Gene expression analysis of shRNA-mediated knockdown of PCID2 in J-Lat 10.6 cells relative to shControl and normalized to cyclophilin A. Bars represent mean of 3 independent shRNA-mediated knockdown experiments and error lines represent SEM. Statistical significance was determined by t-test; ***p<0.001. C. Fold increase in the % of GFP (left y axes) and viability (right y axes) in shControl or shPCID2 J-Lat A2 cells as measured by flow cytometry. Bars represent mean of 3 independent shRNA-mediated knockdown experiments and error lines represent SEM. Statistical significance was determined by t-test; *p<0.05. D. Gene expression analysis of shRNA-mediated knockdown of PCID2 in J-Lat A2 cells relative to shControl and normalized to cyclophilin A. Bars represent mean of 3 independent shRNA-mediated knockdown experiments and error lines represent SEM. Statistical significance was determined by t-test; ***p<0.001.

**Supplementary** Figure 3. **Effect of PCID2 overexpression on HIV-1 gene expression**

**A**. Western blot analysis of PCID2-Flag in transiently co-nucleofected Jurkat cells with pNL4.3.Luc.R-E-, pRL Renilla and either pBud-Control or pBud-PCID2-Flag at 300 or 600 ng. Cells were collected 48 hours post-nucleofection. Beta tubulin was used as a loading control. B. Change in HIV-1 gene expression in Jurkat cells transiently expressing PCID2-Flag was assessed by luciferase assay. Jurkat cells were co-nucleofected pNL4.3.Luc.R-E-, pRL Renilla and pBud-Control or pBud-PCID2-Flag at 300 or 600 ng. Firefly luciferase units were normalized to Renilla luciferase units and relative change in luciferase activity was normalized to control. Bars represent mean of 3 independent experiments and error lines represent SEM. Statistical significance was determined by t-test; ***p<0.001.

**Supplementary** Figure 4. **FISH-Flow analysis of shControl and shPCID2 J-Lat 11.1 cells**

**A**. Representative confocal microscopy images from shControl and shPCID2 J-Lat 11.1 cells seeded as a whole population after FISH-Flow analysis as shown in Figure 5C-E.

**Supplementary** Figure 5. **Mass spectrometry hit list filtering flowchart**

**A**. IP-western blot of PCID2-Flag in control and stably PCID2-Flag expressing J-Lat 11.1 samples for subsequent mass spectrometry analysis. **B**. Filtering scheme used for the hits identified in the two IP-MS runs (left and right columns). Numbers represent total amount of hits after each filtering step. First quadrant shows total hits identified. Total amount of hits were filtered out for common contaminants, including keratin and immunoglobulin proteins. Next, we filtered for unique peptides >1 followed by a fold-enrichment calculation in iBAQ values with a 1.2 fold enrichment cut-off.

**Supplementary** Figure 6. **Efficiency of shRNA-mediated knockdown for TREX2 subunits**

**A**. Gene expression analysis of shRNA-mediated knockdown and GFP mRNA fold induction in shENY2 (A), shCETN3 (B), shDSS1 (C), shMCM3AP (D) cells relative to shControl and normalized to cyclophilin A. Bars represent mean of three independent shRNA-mediated knockdown experiments and error lines represent SEM. Statistical significance was determined by t-test; *p<0.05, **p<0.01, ***p<0.001.

