## Supplementary files for "PCID2 dysregulates transcription and viral RNA processing to promote HIV-1 latency"

### Supplementary Figure 1

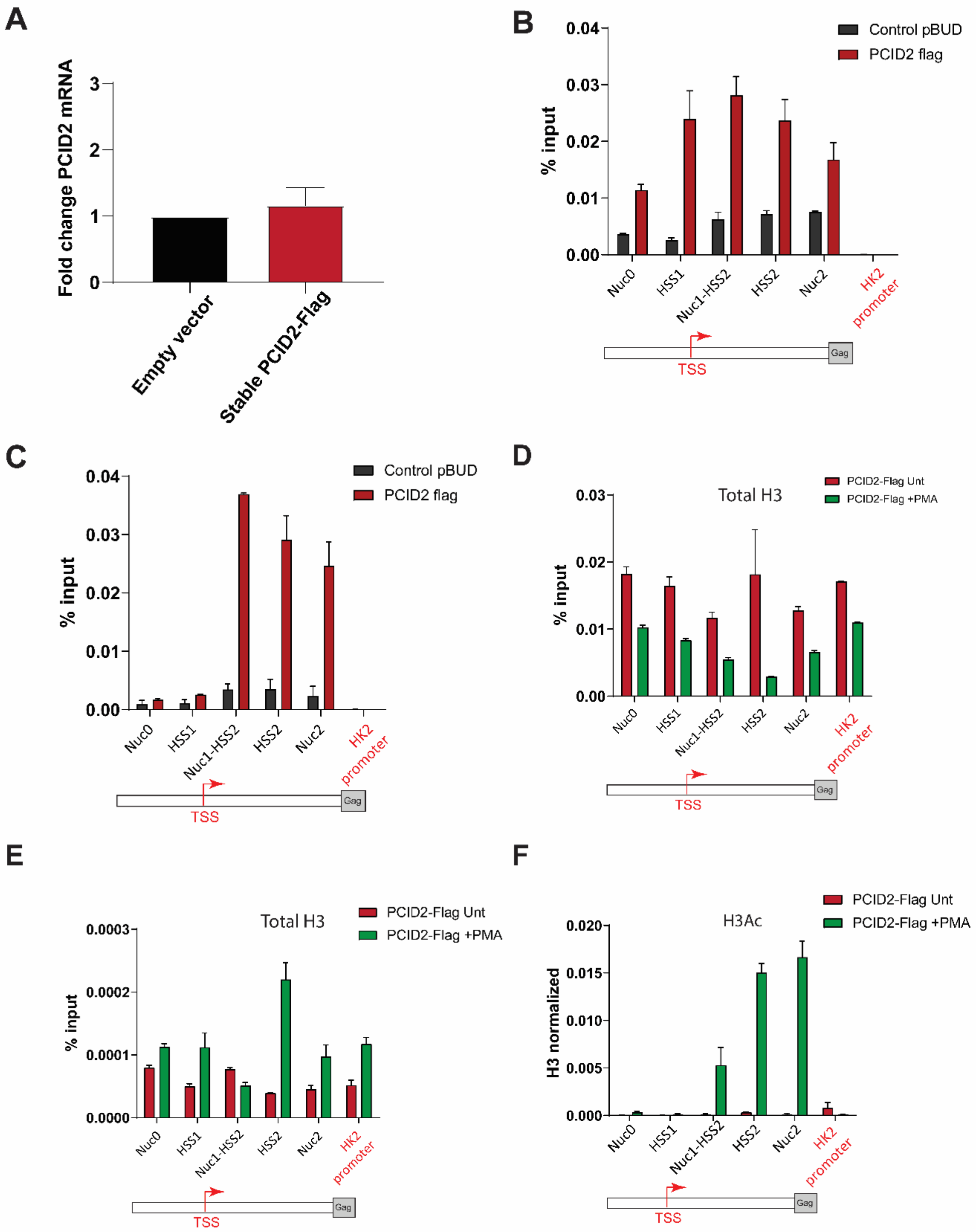

### Supplementary Figure 2

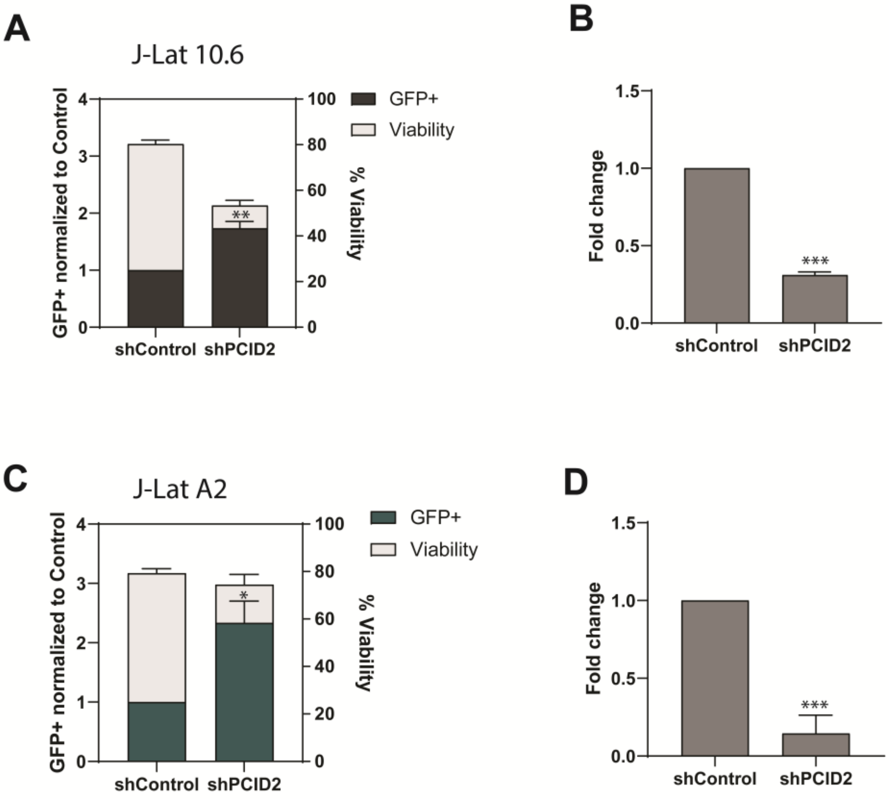

### Supplementary Figure 3

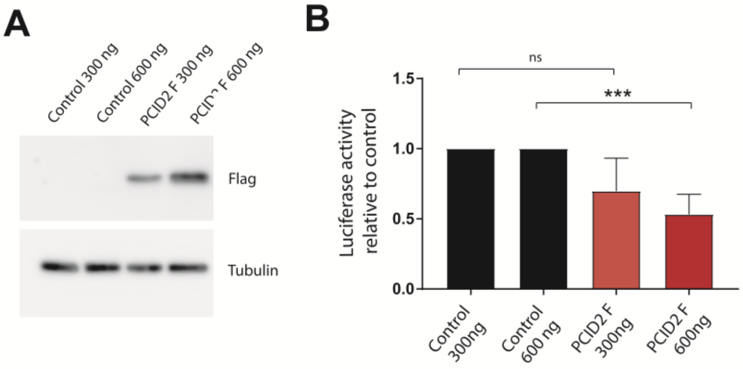

### Supplementary Figure 4

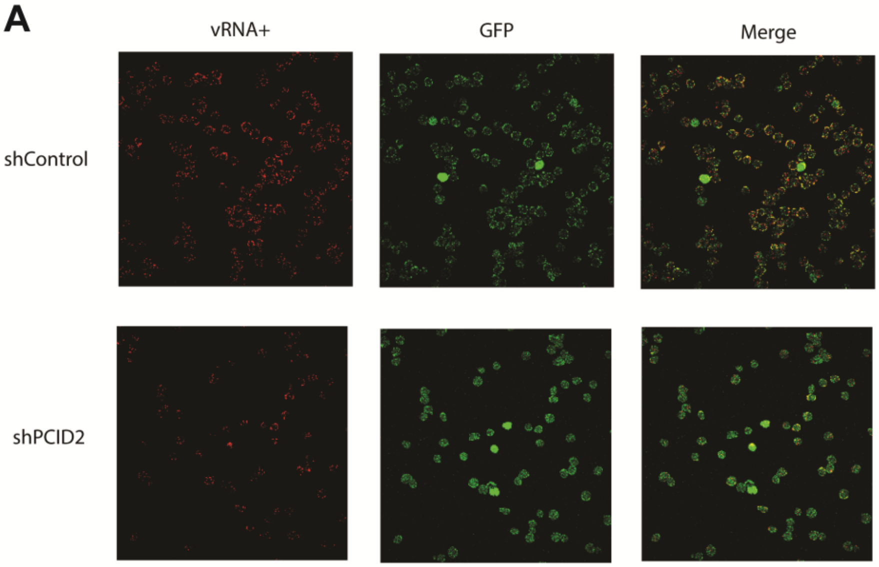

### Supplementary Figure 5

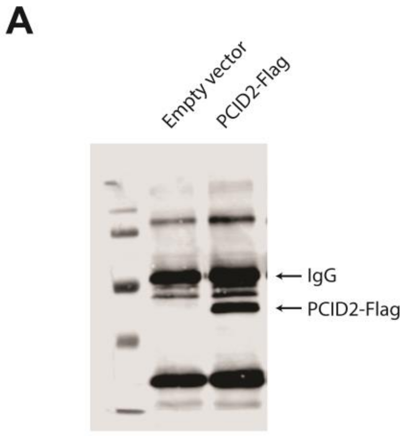


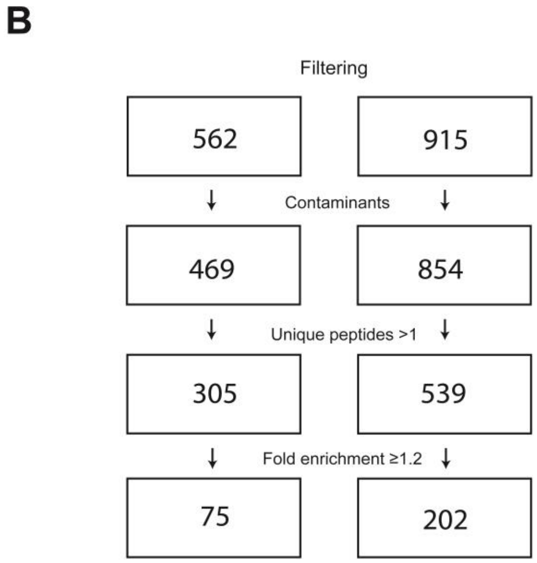

### Supplementary Figure 6

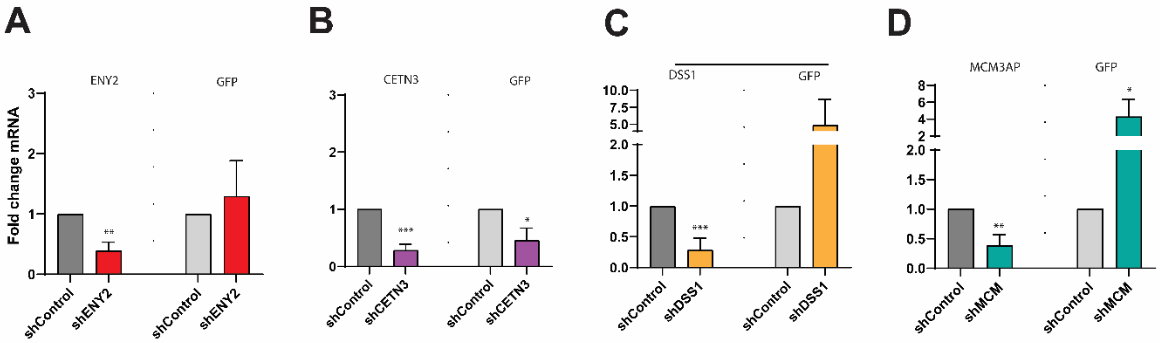
